# Sec17/Sec18 can support membrane fusion without help from completion of SNARE zippering

**DOI:** 10.1101/2021.02.16.431494

**Authors:** Hongki Song, Thomas Torng, Amy Orr, Axel T. Brunger, William Wickner

## Abstract

Membrane fusion requires R-, Qa-, Qb-, and Qc-family SNAREs that zipper into RQaQbQc coiled coils, driven by the sequestration of apolar amino acids. Zippering has been thought to provide all the force driving fusion. Sec17/SNAP can form an oligomeric assembly with SNAREs with the Sec17 C-terminus bound to Sec18/NSF, the central region bound to SNAREs, and a crucial apolar loop near the N-terminus poised to insert into membranes. Though Sec17 aids zippering, we now report that Sec17 and Sec18 will drive robust fusion without requiring zippering completion. Zippering-driven fusion is blocked by deleting the C-terminal quarter of any Q-SNARE domain or by replacing the apolar amino acids of the Qa-SNARE which face the center of the 4-SNARE coiled coils with polar residues. These blocks to fusion, singly or combined, are bypassed by Sec17 and Sec18, and SNARE-dependent fusion is restored without help from completing zippering.

## Introduction

Membrane fusion is central to intracellular protein delivery, neurotransmission, hormone secretion, and cell growth. Its fundamental mechanisms are conserved among organelles and from yeast to humans. Organelle-specific Rab-family GTPases bind tethering proteins to hold membranes together. SNARE proteins have a central role in fusion. SNAREs are classified in 4 families, termed R, Qa, Qb, and Qc (Fasshauer et al., 1998). Each has an N-domain, an α-helical SNARE domain of 50-60 aminoacyl residues with heptad-repeat apolar residues, and often a C-terminal membrane anchor. Each SNARE α-helical turn is termed a “layer.” The central “0-layer” of each fully-assembled SNARE complex has inwardly-oriented arginyl (for R-SNAREs) or glutaminyl (for Qa, Qb, and Qc SNAREs) residues, forming a polar center to the otherwise hydrophobic core of the 4-helical SNARE bundle (Sutton et al., 1998). The SNARE domain layers are numbered from the 0-layer, in the positive direction toward the SNARE C-termini and their membrane anchors and in the negative direction toward the SNARE N-domains. Prior to assembly of 4-SNARE complex, the individual SNARE domains are substantially random-coil (Fasshauer et al., 1997; Hazzard et al., 1999). After membrane tethering by Rabs and effectors, Sec1/Munc18 (SM) family proteins catalyze the N to C directional assembly of SNAREs anchored to each tethered membrane (Fiebig et al., 1999; Sorensen et al., 2006; Baker et al., 2015; Orr et al., 2017; Jiao et al., 2018) into a *trans* RQaQbQc coiled coils complex. SNARE complex assembly is accompanied by the processive transition of each SNARE domain from random coil to α-helix as the heptad-repeat apolar amino acyl residues become sequestered into the interior of the coiled coils (Fasshauer et al., 1997; Sutton et al., 1998). This hydrophobic collapse relies on the exclusion of water by the packing of the 4 SNARE domain helices and is the driving force for SNARE assembly (Sorensen et al., 2006). Completion of SNARE zippering can promote fusion by releasing ∼40kBT per SNARE complex (Gao et al., 2012; Min et al, 2013; Zhang, 2017) to overcome the estimated 40-90 kBT hydration barrier for membrane stalk formation, the dominant energy barrier for fusion (Aeffner et al., 2012). Upon fusion, the *trans*-SNARE complex becomes a *cis*-complex which is anchored to the fused membrane bilayer. Sec17 (αSNAP) and SNAREs serve as a receptor for the Sec18 (NSF) AAA ATPase (Clary et al., 1990; Winter et al., 2009; Zick et al., 2015). Sec18 uses the energy from ATP binding and hydrolysis to disassemble SNAREs for further fusion cycles (Söllner et al., 1993; Ungermann et al., 1998; Zhao et al., 2015) and to disassemble dead-end SNARE complexes to obtain the most fusogenic complexes (Xu et al., 2010; Lai et al, 2017; Choi et al., 2018; Song and Wickner, 2019; Jun and Wickner, 2019).

The molecular interactions between Sec18/NSF, Sec17/αSNAP, and neuronal SNAREs were illuminated by determination of their structures when assembled without membrane anchors into a 20s particle (Zhao et al., 2015; White et al., 2018). The heart of these structures is the 4-helical bundle of the R, Qa, Qb, and Qc SNARE domains. Up to 4 Sec17/αSNAP form a right-handed assembly surrounding the left-handed superhelical coiled coils of the SNARE complex. In this structure, the N-terminal apolar loop of each αSNAP is poised to enter a lipid bilayer adjacent to the SNARE transmembrane domains, the acidic lumenal surface of the four assembled Sec17s may form ionic associations with basic surface side-chains of the SNARE coiled coils, and the C-terminal domain of Sec17/αSNAP contacts Sec18/NSF.

Yeast vacuole fusion is a model of non-neuronal intracellular membrane fusion. It has been studied *in vivo* (Wada et al., 1992), *in vitro* with the isolated organelle (Wickner, 2010), and in a reconstituted proteoliposome-based reaction with all purified recombinant proteins and specified lipids (Mima et al., 2008; Stroupe et al., 2009; Zick and Wickner, 2016). Each protein implicated by the *in vivo* genetics is required for the *in vitro* reconstitution. Fusion requires the Rab Ypt7, the R-SNARE Nyv1 and Q-SNAREs Vam3, Vti1, and Vam7 (hereafter referred to as simply R, Qa, Qb, and Qc), and a large hexameric protein termed HOPS (**<u>ho</u>**motypic fusion and vacuole **<u>p</u>**rotein **<u>s</u>**orting) with multiple direct affinities. Two of the HOPS subunits bind to Ypt7 anchored on each membrane (Brett et al., 2008), explaining how HOPS and Ypt7 mediate tethering (Hickey and Wickner, 2010). A third HOPS subunit is Vps33, the vacuolar Sec1/Munc18 (SM) protein, with direct capacity to bind R and Qa SNARE domains, in parallel and in register (Baker et al., 2015). HOPS also has direct affinity for the Qb and Qc SNAREs (Stroupe et al., 2006; Song et al., 2020) and for vacuolar lipids (Orr et al., 2014). Some, but not all, of the functions of HOPS correspond to fusion factors in other systems. In Ca^2+^-triggered exocytosis, Munc13 cooperates with Munc18 in SNARE assembly (Richmond et al., 2001; Ma et al., 2011; Lai et al., 2017). Munc18 corresponds to the HOPS subunit Vps33, but the corresponding Munc13 function for vacuole fusion is unclear. The association of HOPS with Ypt7 and vacuolar lipids allosterically activates HOPS as a catalyst of SNARE complex assembly (Torng and Wickner, 2020). When the SNAREs are initially in 4-SNARE complexes on two apposed membranes, fusion requires Sec17, Sec18, and ATP to disassemble these *cis*-SNARE complexes and liberate the SNAREs for assembly into *trans*-complexes prior to fusion (Mayer et al., 1996; Nichols et al., 1997; Zick et al., 2015).

Though early studies of vacuole fusion suggested that this disassembly of *cis*-SNARE complexes between rounds of fusion might be the sole function of Sec17 and Sec18 (Mayer et al., 1996), recent findings have broadened our understanding of their roles. While SNAREs are a core fusion machine (Weber et al., 1998), a complete fusion machine (Mima et al., 2008; Zick et al., 2016) also requires the Rab, its tethering effector, an SM-family protein, and the NSF/Sec18 and αSNAP/Sec17 SNARE chaperones. In this more complete context, Sec17 and Sec18 not only disassemble SNAREs between rounds of fusion but also promote fusion *per se*: **<u>1</u>**. *trans*-SNARE complexes which form in a Ypt7-dependent manner between vacuoles bear Sec17 in comparable abundance to the SNAREs (Xu et al., 2010). **<u>2</u>**. Fusion between proteoliposomes has been reconstituted with purified components. When tethering is by nonspecific agents, fusion is inhibited by Sec17, Sec18, and ATP (Mima et al., 2008; Song et al., 2017). However, Sec17, Sec18, and ATP stimulate fusion with HOPS (Mima et al., 2008; Zick et al., 2015; Song et al., 2017). **<u>3</u>**. A pioneering study by Schwartz and Merz (2009) showed that the Qc3Δ deletion of several heptads at the C-terminus of the Qc SNARE blocks the fusion of isolated yeast vacuoles, but that this block could be overcome by the addition of Sec17. Qc3Δ, ending at the SNARE domain layer +3 and thus lacking layers +4 to +8, blocks vacuole fusion *in vivo* as well, and overexpression of Sec17 partially restores cellular vacuole morphology (Schwartz et al., 2017). Reconstituted *in vitro* fusion with limiting Sec17 concentrations, where Sec17 will not restore fusion, also needs Sec18 (ibid). It has been unclear whether the fusion bypass by Sec17 of the deletion of this C-terminal Qc region is particular to just this one SNARE or is general for any arrest of SNARE zippering, and whether Sec17 simply contributes its SNARE-binding energy to the energy of 3-SNARE zippering or whether it drives fusion by other means. **<u>4</u>**. Fusion reactions in which the SNAREs are initially all separated can require Sec17, and this fusion is stimulated by Sec18 without requiring ATP hydrolysis (Zick et al., 2015, Song et al., 2017). Sec17 alone stimulates the fusion of reconstituted proteoliposomes with wild-type SNAREs, and the degree of stimulation is a function of the lipid headgroup and fatty acyl composition (ibid). An intermediate in fusion accumulates during HOPS-dependent fusion without Sec17, allowing a sudden burst of fusion upon Sec17 addition (ibid). At limiting Sec17, fusion was further stimulated by Sec18 without ATP hydrolysis. Single molecule pulling studies have also revealed that αSNAP stabilizes SNARE complexes (Ma et al., 2016) and that it may facilitate their assembly. However, the mechanism whereby HOPS-dependent fusion is stimulated by Sec17/Sec18 has been unclear.

The folding energy of SNARE zippering is considered essential for SNARE-dependent membrane fusion (Sorensen et al., 2006). While Sec17 will promote this zippering, we now report that it has a second mode of promoting fusion which is independent of the energy of zippering. Fusion with SNAREs and HOPS is completely arrested when several heptads repeats in the C-terminal region of the SNARE core complex are removed from any one of the three Q-SNAREs. With any such C-terminally truncated Q-SNARE domain, or even with C-terminal truncations to both Qb and Qc, this block to fusion is bypassed by Sec17 and Sec18 without needing ATP hydrolysis. Association between the C-terminal heptads of the R and the single remaining full-length Qa-SNARE, each anchored to one of the docked membranes, would not yield the same folding energy as with wild-type SNAREs (Sutton et al., 1998) and would contribute far less force towards the bilayer rearrangements of fusion. The N-terminal apolar loop of Sec17 is particularly important for this function of Sec17, and may stabilize SNARE bundles or trigger fusion by insertion into lipid bilayers. Zippering-driven fusion is also arrested with full-length SNARE domains when the apolar, inward-facing residues of the Qa SNARE layers +4 to +8 are replaced by Ala, by Ser, or by Gly, but in each case fusion is restored by Sec17, Sec18, and nonhydrolyzable ATPγS. Strikingly, even fusion which is blocked by the concurrent replacement of apolar residues from the +4 to +8 layers of Qa and the deletion of the +4 to +8 layers of both Qb and Qc, removing all capacity for hydrophobic collapse between the +4 and +8 layers of the R and Q SNAREs, is fully restored by Sec17 and Sec18. We propose that Sec17 either juxtaposes SNAREs through entropic confinement of the remaining full-length SNARE domains or directly promotes bilayer remodeling through insertion of the apolar loops of several SNARE-bound Sec17s, or acts by a combination of these two mechanisms.

## Results

Vacuole SNAREs (Figure 1A) have an N-domain and a SNARE domain. Several have a transmembrane (TM) anchor, but Qc lacks a hydrophobic membrane anchor, and instead associates with membranes through its affinities for the other SNAREs, for HOPS (Stroupe et al., 2006), and for phosphatidylinositol 3-phosphate through its N-terminal PX domain (Cheever et al., 2001). The layers of each SNARE domain are conventionally numbered in both directions from the central, polar 0-layer as shown. Fusion requires that R- and at least one Q-SNARE be anchored to docked membranes (Song and Wickner, 2017). When they are, soluble forms of the other Q-SNAREs without membrane anchors, termed sQ (Figure 1A), will support Ypt7/HOPS-dependent fusion. Based on the EM structure of the homologous human neuronal NSF/αSNAP/SNARE complex (Zhao et al., 2015), we modelled the associations of Sec17 (αSNAP) (Figure 1B) and Sec18 (NSF) with vacuolar SNAREs, viewed in profile or in an end-on view from the membrane (Figure 1C). In this model, we assume that four Sec17 monomers (Figure 1B) assemble together, surrounding the 4-SNARE coiled coil (Figure 1C). In contrast, only two αSNAP molecules have been observed in EM structures of the NSF/αSNAP/V7-SNARE complex (Zhao et al., 2015) and the NSF/αSNAP/SNARE complex that included the linker between the two SNAP-25 SNARE domains (White et al., 2018). The presence of the SNAP-25 linker in these two complexes interferes with binding of the other two αSNAP molecules. Since the vacuolar SNARE complex does not contain a linker between SNARE domains, it is reasonable to postulate that four Sec17 molecules bind to the vacuolar SNARE complex, but the precise number of Sec17 molecules is yet to be determined for the Sec18/Sec17/vacuolar SNARE complex. Rapid fusion needs Sec17 and Sec18 in addition to HOPS and SNAREs (Song et al., 2017). To understand how they work together, we first established that shortened SNAREs will form stable assemblies, then exploited direct assays of SNARE associations to show that HOPS allows Sec17 to promote completion of zippering and SNARE complex stability, and finally examined their functional relationships to show that SNARE-bound Sec17 and Sec18 can promote rapid fusion when energy from zippering is greatly reduced or lost.

**Figure 1.**
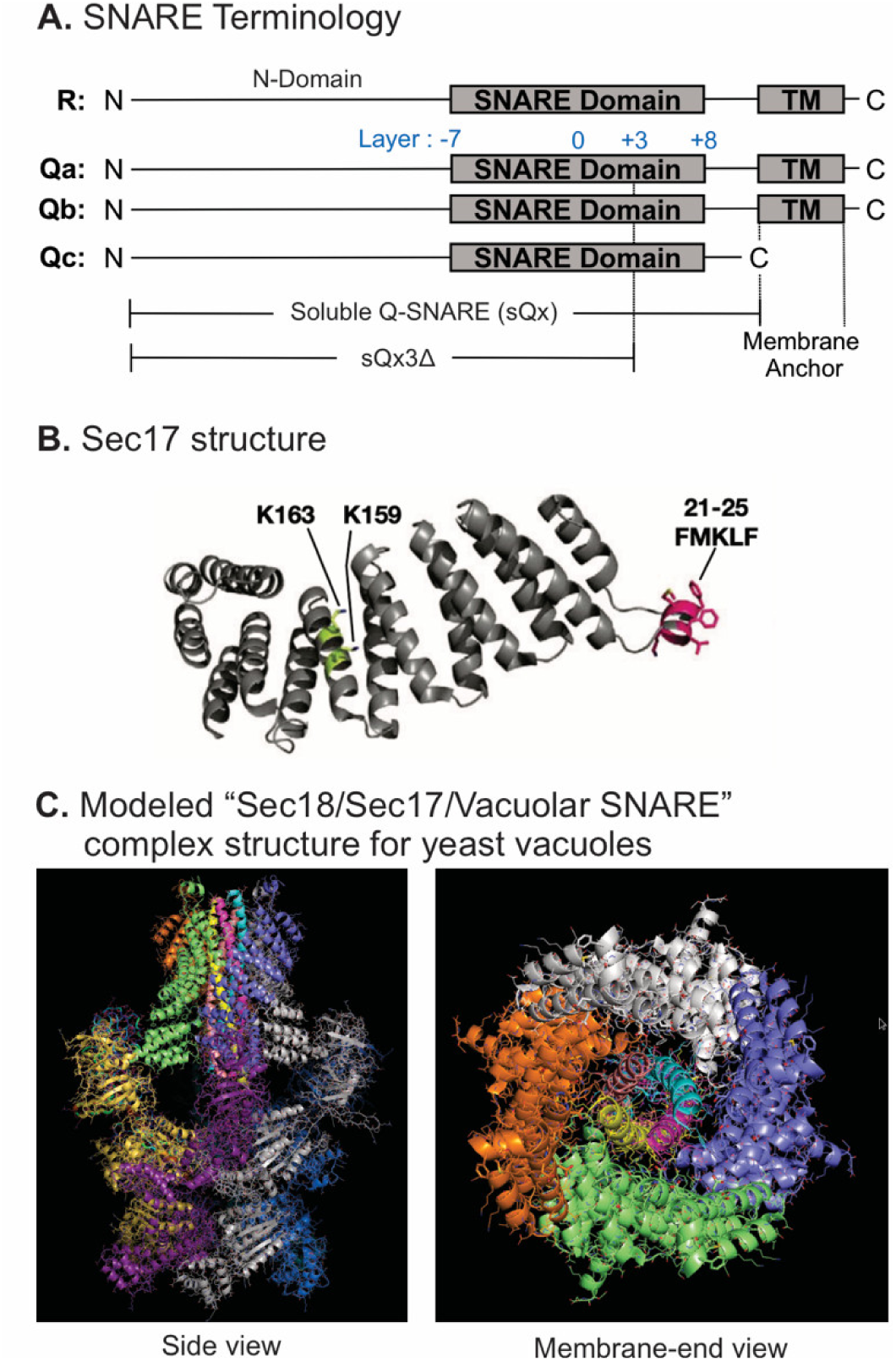
Model of the Sec18/Sec17/vacuolar SNARE complex and Sec17 mutations. **A**. Schematic of the 4 yeast vacuolar SNAREs, the soluble Q-SNAREs (sQx), and their deletion derivatives sQx3Δ lacking regions C-terminal to the +3 layer of the SNARE domain. **B**. Structure of Sec17 (Rice and Brunger, 1999). Residues mutated in certain experiments are highlighted. **C**. Modelling of the vacuolar Sec18/Sec17/SNARE complex. MODELLER (Webb & Sali, 2016) was used to create individual homology models of the vacuolar SNARE complex (Nyv1, Vam3, Vti1, Vam7), and of Sec18 starting from the coordinates of synaptobrevin-2, SNAP-25, syntaxin-1A and NSF in the Cryo-EM structure of the neuronal 20S complex (PDB ID 3J96) (Zhao et al., 2015). These homology models, together with the crystal structure of Sec17 (PDB ID 1QQE) (Rice & Brunger, 1999) were fit into the Cryo-EM structure of the neuronal 20S complex (PDB ID 3J96) (Zhao et al., 2015). We used PDB ID 3J96 because this Cryo-EM structure did not include the SNAP-25 linker and the Habc domain of syntaxin-1A. The vacuolar SNARE complex (Nyv1, Vam3, Vti1, Vam7) also does not include a linker between any of the SNARE motifs; in all structures of NSF/αSNAP/ternary SNARE complexes determined thus far, four αSNAP molecules are observed for SNARE complexes that do not contain a linker connecting two of the SNARE domains (White, Zhao, Choi, Pfuetzner, & Brunger, 2018), and we therefore included four Sec17 molecules in our model of the vacuolar 20S complex. In the PDB coordinate file supplied as Source Data File of the homology model of the Sec18/Sec17/vacuolar SNARE complex, Sec18 molecules, chains A -F; Sec17, chains G-J; Nyv1, chain K; Vam3, chain L; Vti1, chain M; Vam7, chain N. Colors: cyan: Nyv1; magenta: Vam3; yellow: Vti1; salmon: Vam7; gray, orange, green, slate: Sec17; yellow, magenta, gray, blue, salmon, green: Sec18. Cartoon representations are shown. Two views related by a 90 degree rotation are shown (left: sideview; right: membrane-end view).

### SNARE associations during partial zippering

Purified full-length vacuolar SNAREs will spontaneously, though slowly, assemble into a parallel 4-helical coiled coils structure which extends from the N-terminal (-7) layer of the SNARE domain to the C-terminal (+8) end of the SNARE domain. While the association of neuronal SNAREs has been studied extensively, we asked whether vacuolar SNAREs with C-terminal truncations, corresponding to partial zippering, would form stable complexes. Recombinant full-length R-SNARE, with a 6his-tag on its N-terminus, was mixed in octylglucoside with full-length Q-SNAREs, with soluble Q-SNAREs (sQ), or with sQ3Δs. Stable spontaneous complexes were isolated with nickel-NTA beads and examined by immunoblot (Figure 2, and Figure 2-supplement 1). Complex formation with full-length SNAREs (lane 1) was not blocked by the absence of trans-membrane domain from Qa, Qb, or both (lanes 2-4) or by the 3Δ deletion in the C-terminal end of the SNARE domain in any one or two Q-SNAREs (lanes 5-10). This capacity of vacuolar SNAREs to form stable partially-zippered structures raised the question of whether these SNAREs can zipper efficiently, especially when anchored to membranes or associated with other fusion proteins such as HOPS.

**Figure 2.**
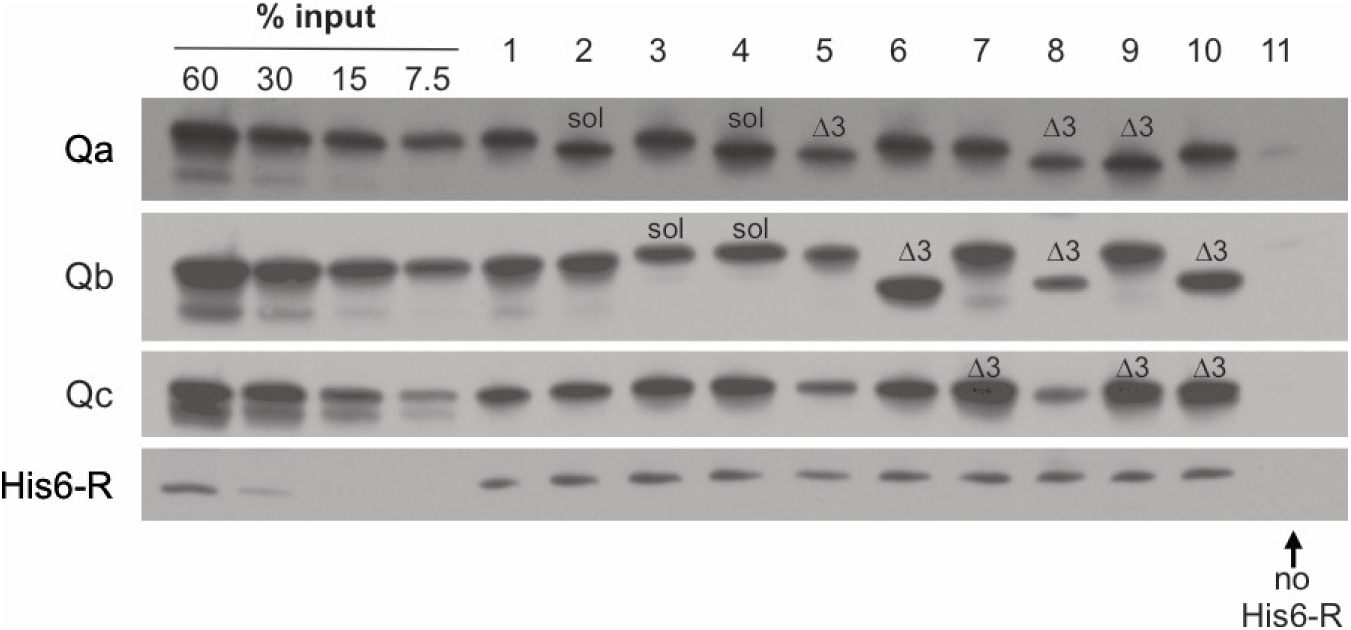
C-terminal truncations allow stable 4-SNARE complex assembly. Reactions (50µl) containing 2.5µM of each SNARE in Buffer 151 (20mM HEPES-NaOH pH 7.4, 150mM NaCl, 10% glycerol, 1% ß-OG, 20mM imidazole) were nutated at 4°C for 2 hours. A 40µl portion of each was transferred to a 0.5ml Eppendorf tube containing 20µl of a 50% slurry of Qiagen Nickel-NTA Agarose in Buffer 151. Suspensions were nutated at 4°C for 1 hour. Loosely bound proteins were removed by four successive suspensions in 0.5ml Buffer 151, each followed by centrifugation (500xg, 6 min, 4°C) and removal of the supernatant. Bound proteins were eluted by boiling the agarose in 50µl of SDS sample buffer (50mM TrisCl, pH 6.8, 5% glycerol, 2% sodium dodecyl sulfate, 0.2% bromophenol blue, 1% ß-mercaptoethanol) for 5 minutes. Samples were analyzed by SDS-PAGE followed by immunoblotting.

### Sec17-induced completion of zippering

To study the kinetics of SNARE interactions, we employed a fluorescence resonance energy transfer (FRET) assay of vacuolar SNARE associations (Torng and Wickner, 2020). The Qc-SNARE was prepared with a unique cysteinyl residue in any of 3 positions (Figure 3A), either the native Cys208 which is upstream (U) of the SNARE domain or, after substitution of serine for this cysteine, with Met250Cys at the N-terminal end of the SNARE domain or with Ala316Cys at the C-terminal end of the SNARE domain. Each was derivatized with Oregon Green 488. Qb-SNARE domain fusion proteins were also prepared with a unique cysteine either immediately N-terminal, or C-terminal, to the Qb SNARE domain, and each of these was derivatized with Alexa Fluor 568. These fluorescent proteins were co-incubated with proteoliposomes bearing Ypt7, R and Qa. SNARE complex assembly can occur spontaneously on these proteoliposomes, but assembly is stimulated by HOPS, allowing direct comparison of HOPS-dependent and HOPS-independent SNARE assembly (Torng and Wickner, 2020). SNARE complex assembly with HOPS gave a high average FRET efficiency when fluorophores were at the N-terminus of the Qb SNARE domain and at or near the N-terminal end of the Qc SNARE domain (Figure 3B, red and yellow curves). There was lower average FRET efficiency when the fluorophores were at opposite ends of the Qc and Qb SNARE domains (Figure 3B, blue, green, and indigo). When both fluorophores were at the C-terminal ends of the Qb and Qc SNARE domains, a low signal was seen (Figure 3B, purple), similar to the average FRET efficiency when fluorophores were at opposite ends of the SNARE domains and thus suggesting incomplete zippering. After 1 hour, Sec17 was added to each reaction. Strikingly, Sec17 only enhanced the average FRET efficiency between C-terminal fluorophores, rapidly rising to the level seen when the fluorophores were together at the N-terminii of the SNARE domains (Figure 3B, purple; see also Figure 3-figure supplement 1), indicating Sec17-induced zippering. Sec17/α-SNAP has been shown to promote the zippering of isolated SNARE domains (Ma et al., 2016), but is now seen to do so in the context of membranes and the Rab-activated SNARE assembly catalyst HOPS.

**Figure 3.**
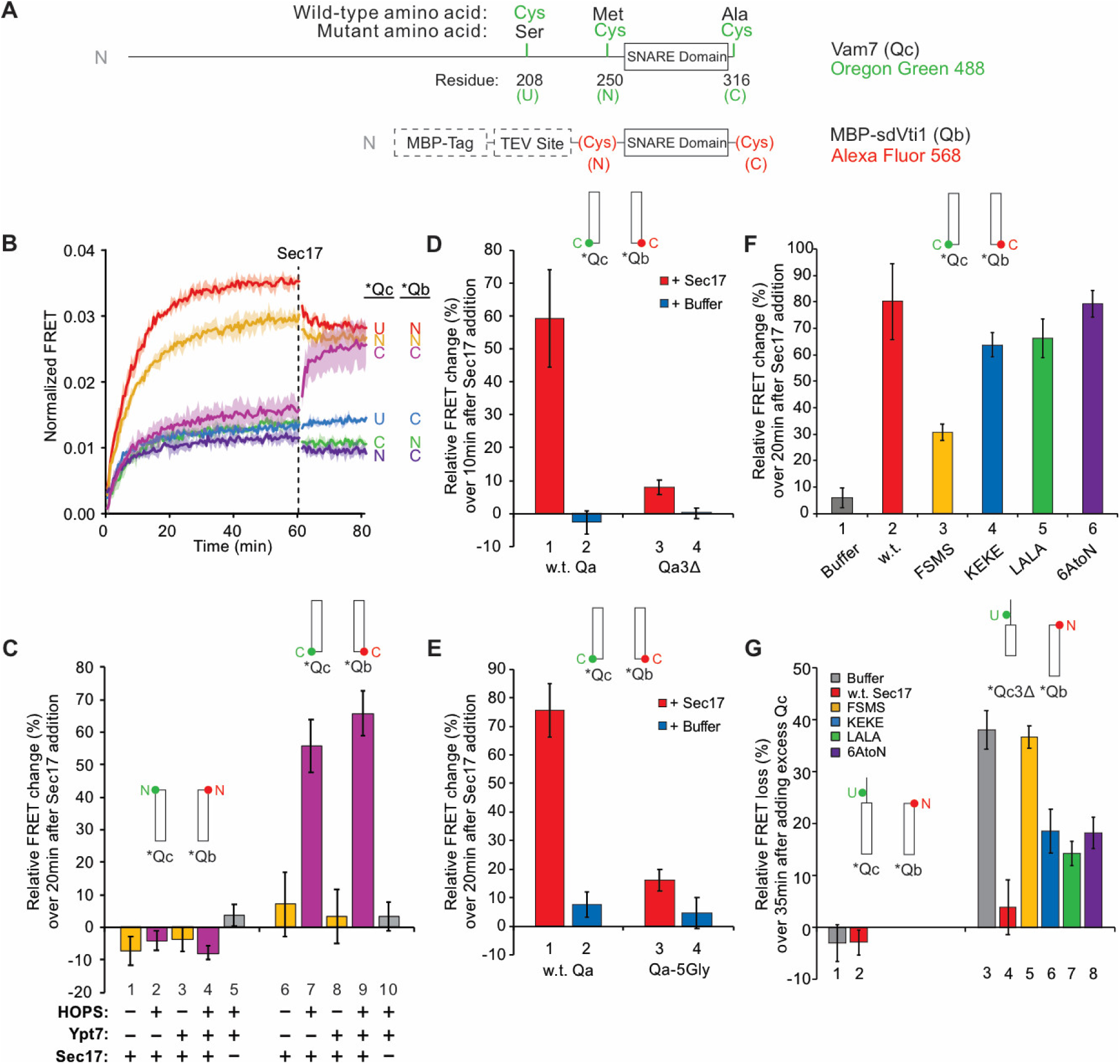
Sec17 promotes complete zippering of full-length SNARE domains in a HOPS-dependent manner and stabilizes complexes with truncated SNARE domains. **A**. Schematic of fluorescently labeled SNARE constructs used in this study. SNAREs were derivatized as described previously (Torng et al., 2020). Wild-type Qc contains a single Cys residue at 208 at the upstream (U) position. (N)- and (C)-labeled constructs replace residues 250 and 316 with Cys, respectively, while also replacing Cys208 with Ser. Each Qc construct was derivatized with Oregon Green 488 as described (Torng et al., 2020). A fusion of maltose binding protein (MBP), a TEV site, and the Qb SNARE domain (residues 133-187) was expressed with an added cysteinyl residue immediately upstream or downstream of the SNARE domain. Each Qb construct was derivatized with Alexa Fluor 568. Qc and Qb labeled with a fluorescent probe at any position are written as *Qc and *Qb. **B-G**. Bar graphs are reported as the mean of the relative FRET change (%) per trial with propagated standard deviation for n=3 trials. The relative change was calculated from averaging 10 data points each from just before Sec17 addition and from the end of the measurement period 20 min later, except where indicated. See Supplementary Data for specific time points used as well as the statistics for the propagation of uncertainty. Also see Supplementary Figure for a bar graph representation for (B) and for the kinetic curves for (C-G). **B**. Sec17 promotes zippering of the C-terminus of the SNARE complex. Ypt7/RQa proteoliposomes were incubated with pairs of *Qb and *Qc labeled at the N-, C-, or upstream (U)- locations as indicated in A. Curves are averages of n=3 trials, and the shaded regions behind each curve show the standard deviation per time point. **C**. Sec17-promoted zippering requires HOPS. RQa proteoliposomes were incubated with either the N-labeled *Qb/*Qc pair (left) or the C-labeled *Qb/*Qc pair (right), with Ypt7 and HOPS as indicated. A reaction with buffer (RB150) added instead of Sec17 serves as a negative control. **D**. Sec17 does not promote C-terminal zippering if the SNARE domain of Qa is truncated. Ypt7/R proteoliposomes were incubated with C-terminally labeled *Qb and *Qc and either soluble Qa or Qa3Δ, and the relative FRET change was calculated over 10 min after Sec17 or buffer addition. **E**. Sec17-induced zippering requires the apolar heptad repeat amino acyl residues in Qa SNARE domain layers +4 to +8. Proteoliposomes bearing Ypt7, R-SNARE, and either wild-type Qa or Qa with the +4 - +8 layers inwardly-oriented apolar residues mutated to Gly were incubated with C-terminally labeled *Qb and *Qc, and then Sec17 or its mutants were added at t=60 min. **F**. Sec17-driven zippering is stunted by the FSMS mutation. Ypt/R proteoliposomes were incubated with sQa and C-terminally labeled *Qb and *Qc, then Sec17 or mutants as indicated were added at t = 60 min. **G**. Sec17 stabilizes incompletely-zippered SNARE complexes. Ypt7/R proteoliposomes were incubated with sQa and C-terminally-labeled *Qb and *Qc. Sec17 or the indicated mutants were added at t = 60 min. Non-fluorescent Qc (8.5 µM) was added at t = 80 min, and the loss of FRET over 35 min was measured starting immediately after addition of non-fluorescent Qc.

### Sec17-induced completion of zippering requires HOPS

Proteoliposomes with R, Qa, and Ypt7 (where indicated) were incubated with Qb-SNARE domain and Qc. Both SNARE domains were either fluorescently labeled at their N-termini (Figure 3C, bars 1-5) or at their C-termini (Figure 3C, 6-10). Incubations were performed in the presence or absence of HOPS. Sec17 addition after hour did not enhance the average FRET efficiency between N-terminally disposed fluorophores in the presence or absence of HOPS (Figure 3C, bars 1-5), but stimulated the average FRET efficiency between SNARE domain C-terminal fluorophores in a HOPS-dependent manner (Figure 3C, bars 6-10). This reflected HOPS-dependent and Sec17-induced full SNARE zippering, as it was greatly diminished when zippering was inhibited by the absence of the +4 to +8 layers of the sQa-SNARE (Figure 3D, bar 1 vs 3) or by the conversion of each inward-facing apolar residue of the full-length Qa SNARE domain layers +4 to +8 to Gly (Figure 3E, bar 1 vs 3). The F22SM23S mutation of Sec17 (FSMS hereafter), diminishing the hydrophobicity of its N-domain loop (Song et al., 2017), reduced the Sec17 capacity for inducing HOPS-dependent zippering (Figure 3F). We also examined the effects of several other mutants of Sec17. The K159E,K163E mutation (KEKE hereafter) diminishes Sec17:SNARE association (Marz et al., 2003); one of these residues (Sec17 K159) is in a pair (αSNAP K122, K163) that abolishes disassembly of the neuronal SNARE complex by NSF/αSNAP (Zhao et al., 2015). The C-terminal L291A,L292A mutation of Sec17 (LALA hereafter) interferes with its cooperation with Sec18 for SNARE complex disassembly (Barnard et al., 1997; Schwartz and Merz, 2009; Zick et al., 2015), and 6AtoN is the conversion of 6 inward-facing acidic residues of Sec17 which face basic SNARE residues in the 20s structure (Figure 1C) to neutral residues. Neither KEKE, LALA, nor 6AtoN had a large effect on the capacity of Sec17 to promote HOPS-dependent zippering (Figure 3F).

Sec17 can also be shown to interact with partially-zippered SNARE complex by its effects on SNARE complex stability. SNARE complex was assembled by HOPS on Ypt7/R proteoliposomes with soluble Qa, with the Qb SNARE domain labeled with a fluorophore at a cysteinyl residue upstream of the SNARE domain, and with Qc of full length (w.t.) or with the 3Δ C-terminal truncation, each bearing a fluorophore at its native cysteinyl residue upstream of the SNARE domain. When the complex of proteoliposomes with these fluorescent Qb and Qc had full-length SNARE domains, it was stable whether or not it included Sec17, as there was no loss of average FRET efficiency after addition of a molar excess of nonfluorescent Qc (Figure 3G, bars 1 and 2; Figure 3-figure supplement 1G). In contrast, fluorescent Qc3Δ was “chased” by exchange with nonfluorescent Qc (bar 3), but Sec17 stabilized this SNARE complex, blocking the chase (bar 4). Thus, in the absence of Sec17, the assembly of Qc3Δ into partially-zippered SNARE complex is reversible. Each domain of Sec17 helps to stabilize Qc3Δ against exchange (bars 5-8), but especially the Sec17 N-terminal apolar loop (bar 5).

### Direct binding of Sec17 to HOPS

Proteoliposome fusion promoted by synthetic tethers is inhibited by all concentrations of Sec17 whereas HOPS-dependent fusion is only inhibited by very high Sec17 levels (Song and Wickner, 2019) and Sec17-triggered zippering requires HOPS (Figure 3C). We therefore tested whether HOPS might directly associate with Sec17. Phosphatidylcholine liposomes were prepared without protein (“naked”) or with TM-Sec17, a recombinant form of Sec17 with an N-terminal apolar transmembrane anchor (Song et al., 2017). These liposomes were incubated with 3 independent preparations of HOPS, then floated through a density gradient. HOPS only bound in the presence of anchored Sec17 (Figure 4 and Figure 4-figure supplement 1), showing a direct and nearly stoichiometric binding between these proteins.

**Figure 4.**
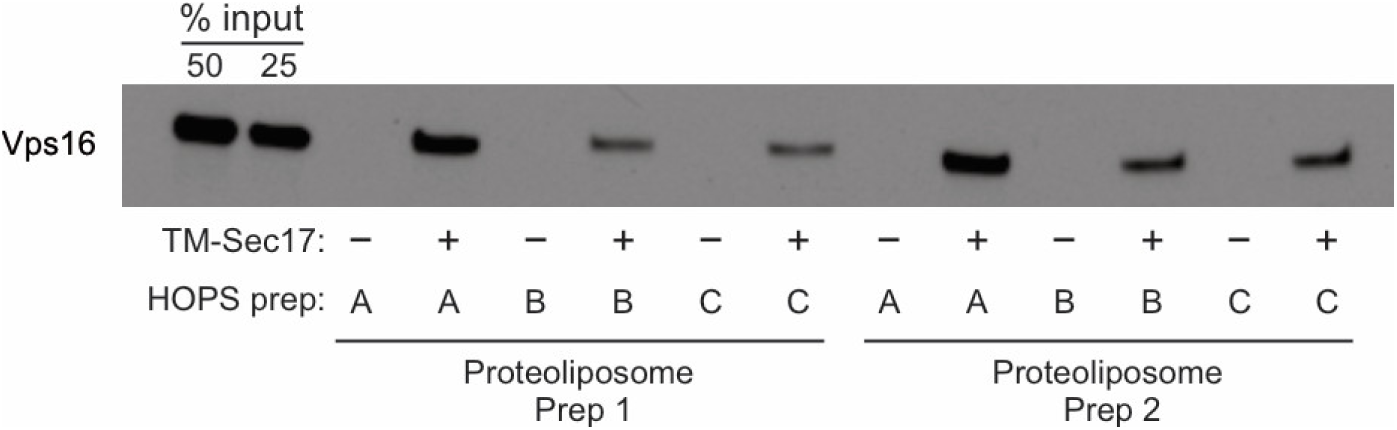
HOPS binds Sec17. **A**. Liposomes composed of PC/PS/NBD (83.5/15/1.5%) were prepared from mixed micellar solutions by detergent dialysis as described in Materials and Methods. Liposomes bore either no proteins or TM-Sec17 (Song et al 2017) at a 1:4000 protein to lipid molar ratio. Floatation assays were conducted as described in Song et al, 2020, Figure 2. In brief, 30μl reactions in RB150 contained 0.5mM liposomal lipid, 125nM integral TM-Sec17 where present, 0.2% defatted BSA, 1mM MgCl2, and 500nM HOPS. Reactions were incubated (1hr, 30°C) and liposomes floated through Histodenz (Sigma, St. Louis, MO) density medium. Collected samples were solubilized in 0.125% Thesit, adjusted for lipid recovered and analyzed for bound proteins by immunoblot for Vps16, a HOPS subunit. Because the reaction contained four times more HOPS than integrally bound TM-Sec17, we express the standard curve as the per cent of molar equivalence (to total liposomal TM-Sec17) of HOPS. **B**. Western blots from three repetitions of this experiment were analyzed by UN-SCAN-IT Software with standard curves. Mean values and standard deviations are shown.

### Generality of Sec17 and Sec18 bypass of arrested zippering

Sec17 can restore fusion when Qc has truncations at the C-terminal end of its SNARE domain (Schwartz and Merz, 2009), stimulated by Sec18 (Schwartz et al, 2017). Since HOPS and Sec17 support SNARE zippering (Figure 3), and vacuolar SNAREs with incomplete SNARE domains can form stable complexes corresponding to incomplete zippering (Figure 2), we asked whether Sec17 restoration of fusion was limited to Qc C-terminal SNARE domain truncations or was seen with similar deletion of residues after the +3 layer in the other Q-SNAREs as well. Reconstituted proteoliposomes bearing Ypt7 and R-SNARE were assayed for fusion with proteoliposomes bearing this Rab and any two anchored Q-SNAREs. Proteoliposome mixtures were incubated with HOPS and the soluble forms of the remaining soluble Q-SNARE deprived of its membrane anchor and a C-terminal portion of its SNARE domain (Figure 5: A, QcΔ3; B, QbΔ3; C, QaΔ3). As reported (Schwartz et al., 2017), there was no fusion with Qc3Δ unless Sec17, Sec18, and ATP or ATPγS were present (Figure 5A). While ATP and its nonhydrolyzable analog ATPγS support comparable fusion with wild-type SNAREs (Song et al., 2017), hydrolyzable ATP inhibits fusion through Sec17/Sec18-mediated disassembly of SNARE complexes when defective SNAREs such as Qc3Δ are present, a proofreading function. The same pattern was seen for fusion with sQb3Δ (Figure 5B) and sQa3Δ (Figure 5C). The unique spatial disposition of the Sec17/αSNAP molecules with respect to each SNARE (Figure 1C, and Zhao et al., 2015) makes it unlikely that Sec17 could somehow fill each of the gaps left by each of these deletions to shield apolar residues within the SNARE bundle and thereby continue to drive zippering, or that Sec17 binding could induce the remaining R and two Q +4 to +8 layers to somehow rotate to form a hydrophobic 2- or 3-layered core.

**Figure 5.**
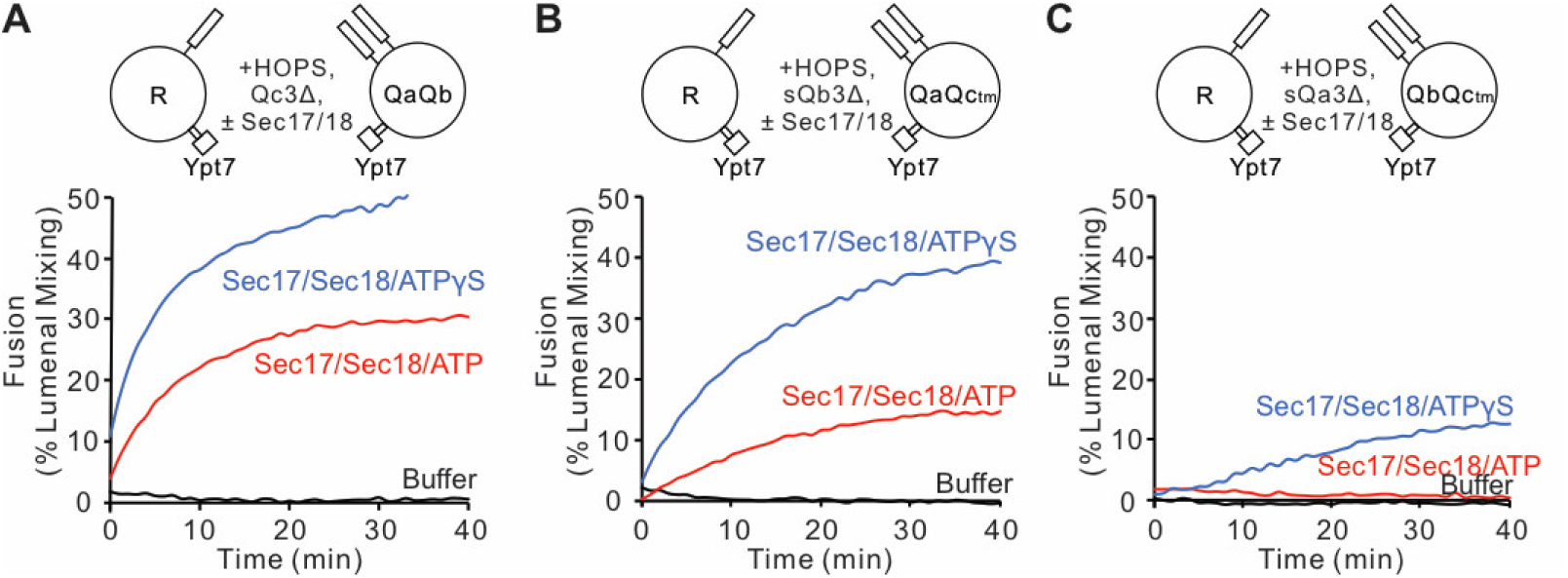
Fusion is blocked by deletion of the last 5 C-terminal SNARE domain layers from any single Q-SNARE and is restored by Sec17, Sec18, and ATP or ATPγS. **A**. Fusion incubations, as described in Methods, had Ypt7/R- and Ypt7/QaQb proteoliposomes (1:8,000 Ypt7:lipid molar ratio, 1:16,000 SNARE:lipid molar ratios), 2μM Qc3Δ, and where indicated 600 nM Sec17, 300 nM Sec18, 1mM ATP (red) or ATPγS (green). **B**. Fusion with 2μM sQb3Δ and with Ypt7/QaQc-tm proteoliposomes, but otherwise as in (A). **C**. Fusion with 2μM sQa3Δ and with Ypt7/QbQc-tm proteoliposomes, but otherwise as in (A).

Fusion assays were also performed with Ypt7/R proteoliposomes and each of the three Ypt7/single anchored Q-SNARE proteoliposomes in the presence of HOPS and the other two soluble Q-SNAREs (Figure 6). With membrane-anchored Qa and with sQb and sQc having complete SNARE domains, there was HOPS-dependent fusion without further addition (Figure 6A, black line), though Sec17 and Sec18 with AMP-PNP, ATPγS, or ATP did stimulate (compare black curves, A-D). Deletion of the +4 to +8 layers from either soluble Qb or Qc abolished fusion (Figure 6A), which was restored by Sec17 and Sec18 with either AMP-PNP, ATPγS, or ATP (B-D, red and blue curves). There was no fusion when both soluble Q-SNAREs had truncated SNARE domains (Figure 6A, orange), but strikingly the fusion was partially restored by Sec17 and Sec18 with ATP (Figure 6D, orange) and more fully restored when the adenine nucleotide was resistant to hydrolysis (Figure 6, B and C, orange). With 2 Q-SNAREs lacking the C-terminal portion of their SNARE domains, the apolar amino acyl residues of the remaining 2 SNAREs would not be as effectively shielded from water if they continued zippering together. Fusion could not be restored by Sec17 and Sec18 if either soluble SNARE was omitted instead of truncated (green and purple). When only the Qb Q-SNARE was membrane anchored (Figure 6E), there was little fusion without Sec17 and Sec18 (Figure 6, E-H, black curves). When the SNARE domain of sQa or Qc had been truncated, fusion was strictly dependent on non-hydrolyzable ATP analogs, and little fusion was seen with dual SNARE domain truncation. Similar patterns were seen with anchored Qc (Figure 6, I-L). We term the fusion induced by Sec17 and Sec18 in the presence of 3Δ SNARE domain truncations “zippering bypass fusion.”

**Figure 6.**
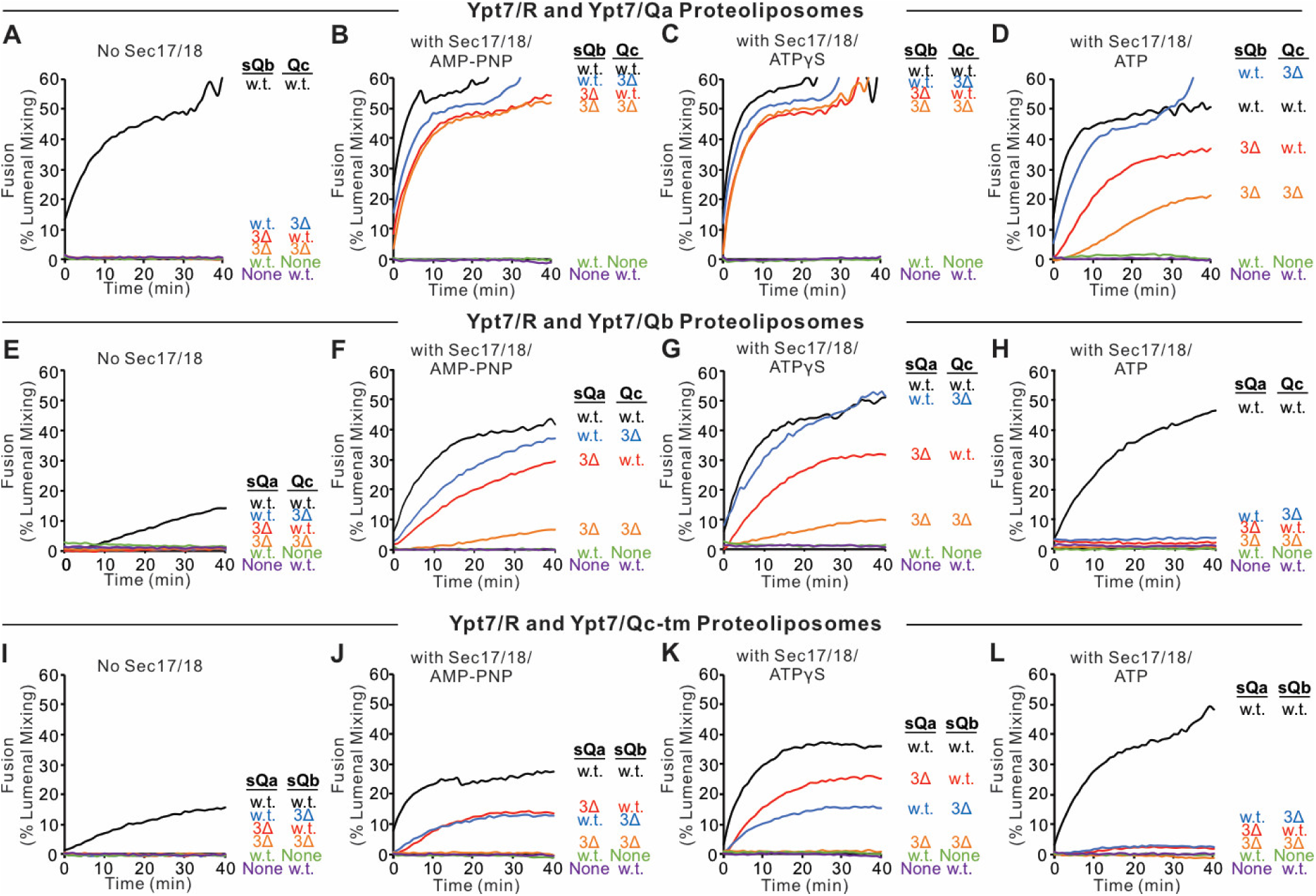
Fusion with single membrane-anchored Q-SNAREs. **A-D**. Fusion incubations, as described in Methods, had Ypt7/R- and Ypt7/Qa-proteoliposomes (1:8,000 Ypt7:lipid molar ratio, 1:16,000 SNARE:lipid molar ratios), 50 nM HOPS, 2μM sQb or sQb3Δ, 2μM Qc or Qc3Δ, and where indicated 600 nM Sec17, 300 nM Sec18, and 1mM ATP, AMP-PNP, or ATPγS. **E-H**. Fusion incubations as for (A), but with Ypt7/Qb-proteoliposomes and 2μM sQa or sQa3Δ, 2μM Qc or Qc3Δ and Sec17, Sec18, and adenine nucleotide as indicated. **I-L**. Fusion incubations as for (A), but with Ypt7/Qc-tm-proteoliposomes and 2μM sQa or sQa3Δ, 2μM Qb or sQb3Δ and Sec17, Sec18, and adenine nucleotide as indicated.

To explore the interactions of Sec17 and Sec18 during zippering-bypass fusion, we assayed the fusion of Ypt7/R and Ypt7/Qa RPLs with sQb3Δ, Qc3Δ, and HOPS, using various concentrations of wild-type or mutant Sec17, and with or without Sec18 and ATPγS (Figure 7). Without Sec17, fusion is not supported by Sec18 (A and B, blue curves). Limited fusion is possible with 1 or 2 μM wild-type Sec17 alone (A, black and red), but Sec18 allows faster fusion and with less Sec17 (A, B; tan). Fusion requires the apolar loop near the N-terminus of Sec17, as the F21S,M22S mutation (FSMS) blocks fusion entirely (C, D). The K159E,K163E mutation (KEKE) diminishes Sec17:SNARE association (Marz et al., 2003). The KEKE mutation prevents fusion without Sec18 (E), but a slow and limited fusion with KEKE-Sec17 is restored by Sec18 (F). The C-terminal L291A,L292A mutation of Sec17 (LALA), which interferes with its cooperation with Sec18 for SNARE complex disassembly (Barnard et al., 1997; Schwartz and Merz, 2009; Zick et al., 2015), diminishes zippering-bypass fusion (red curves, A vs G), and a limited fusion is restored through the addition of Sec18 (H). These data suggest that Sec17 action directly requires its apolar loop domain, since the loss of this apolar region is not bypassed by Sec18. Sec18 may stimulate fusion by either contributing to the binding of Sec17 to *trans*-SNARE complexes, by causing conformational changes in Sec17 or the SNAREs, or both. Basic residues in the +3 to +8 layers near the C-terminus of the R and Qa SNARE domains are near acidic residues on the interior of the Sec17 assembly (Figure 7I). To determine whether Sec17 might rely on these ionic interactions to support fusion, we converted these acidic residues of Sec17 to alanine or serine, creating the mutant Sec17-E34S, E35S, D38S, E73A, D43A, E75A (termed Sec17 6 Acidic to Neutral, or Sec17-6AtoN), but this mutant Sec17 still supports fusion (Figure 7, J and K). Interestingly, mutation of acidic residues of αSNAP near the C-terminal end of the neuronal SNARE complex also only had a modest effect on disassembly activity of NSF/αSNAP (Zhao et al., 2015).

**Figure 7.**
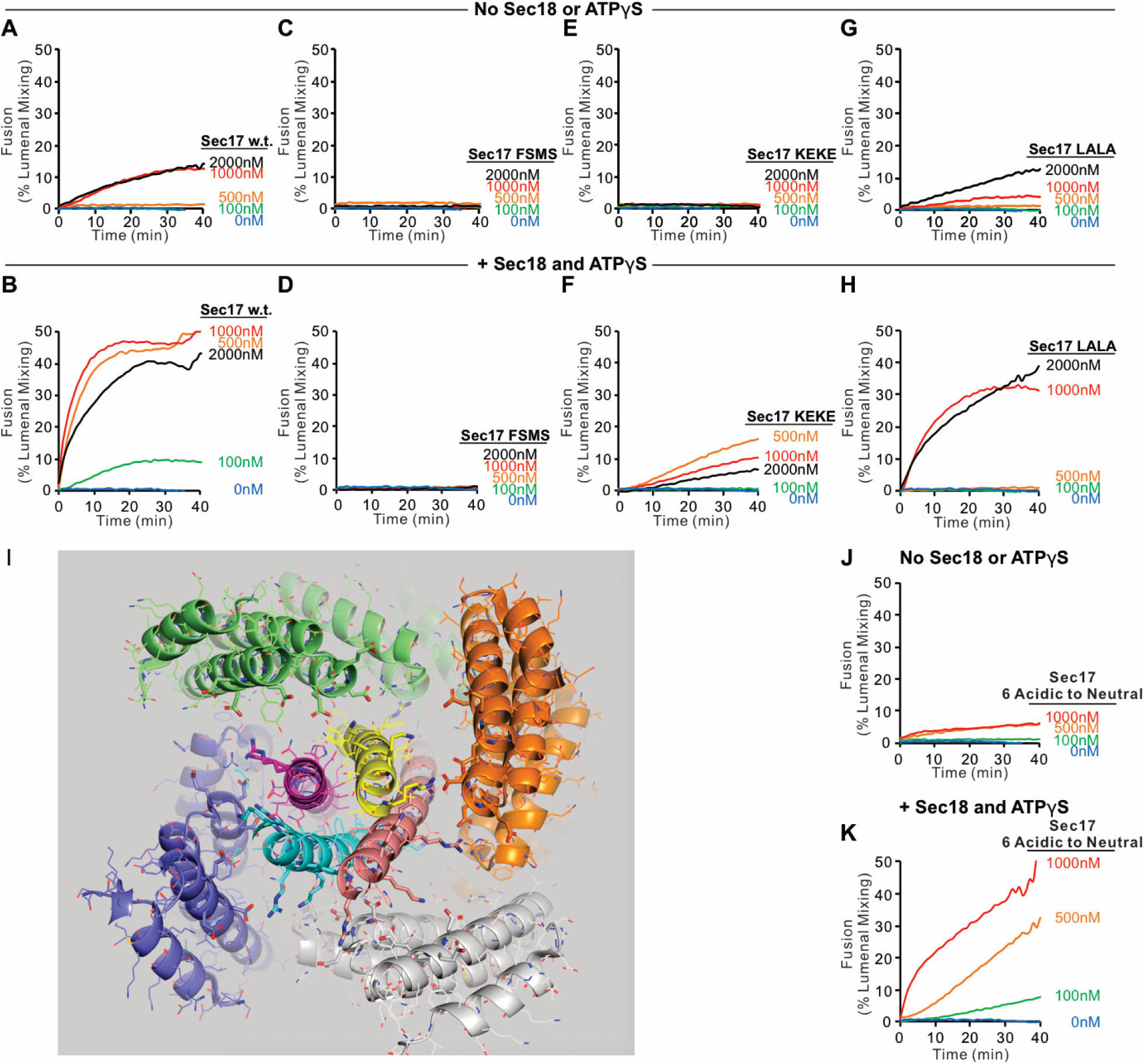
Role of each domain of Sec17 in zippering-bypass fusion. Fusion between Ypt7/R and Ypt7/Qa proteoliposomes (1:8,000 Ypt7:lipid and 1:16,000 SNARE/lipid molar ratios) was assayed with 50 nM HOPS, 2μM sQb3Δ, 2μM Qc3Δ, the indicated concentration of wild-type or mutant Sec17, and with or without 250nM Sec18 and 1mM ATPγS. **I**. Ionic interactions between Sec17s and vacuolar SNAREs in the +4 to +8 layers in the model of the Sec18/Sec17/vacuolar SNARE complex (Figure 1). Colors: cyan: R; magenta: Qa; yellow: Qb; salmon: Qc; gray, orange, green, slate: Sec17; red: oxygen atoms; blue nitrogen atoms. Cartoon representations are shown along with sidechains shown as thin lines. Thick lines (sticks) are interacting glutamate and aspartate (acidic) residues on the surface of Sec17 (aminoacyl residues 34, 35, 38, 73, 74, 75) that interact with the vacuolar SNARE complex lysine and arginine basic residues in each of the 4 SNAREs.

To test whether Sec18 is simply promoting fusion by contributing to the affinity of Sec17 for membranes which bear SNARE complexes, we prepared proteoliposomes with Ypt7, with R or Qa, and with either no Sec17, with an equimolar (to SNAREs) TM-Sec17 which bears an N-terminal hydrophobic transmembrane anchor domain derived from Qb, or with TM-Sec17-F21S,M22S (TM-FSMS hereafter). In the presence of HOPS, sQb3Δ, and Qc3Δ, membrane-anchored Sec17 supported zippering-bypass fusion which was strictly Sec18-dependent (Figure 8, A-C, no Sec18; D-F, with Sec18 and ATPγS). TM-Sec17 on both sets of proteoliposomes supported fusion (Figure 8D and G, black) but fusion was not seen in the absence of TM-Sec17 (Figure 8D and G, red), and only limited fusion was seen with TM-FSMS-Sec17 (8D and G, blue). Fusion was greatly diminished if TM-Sec17 or TM-Sec17-FSMS was only present on one of the fusion partner proteoliposomes (Figure 8E and H), but there was substantial fusion if one of the proteoliposomes had TM-Sec17 and one had TM-Sec17-FSMS (Figure 8F and I). The simplest model is that several Sec17 molecules are required to form the Sec17 assembly around the SNAREs, but that the hydrophobic Sec17 N-domain loop is only necessary on one membrane for fusion. AMP-PNP functioned as well as ATPγS (compare G-I to D-F), but hydrolyzable ATP completely blocked fusion with membrane-anchored TM-Sec17 (J-L), similar to its lower activity with wildtype Sec17 (Figure 6A), reflecting the proofreading activity of Sec17/Sec18.

**Figure 8.**
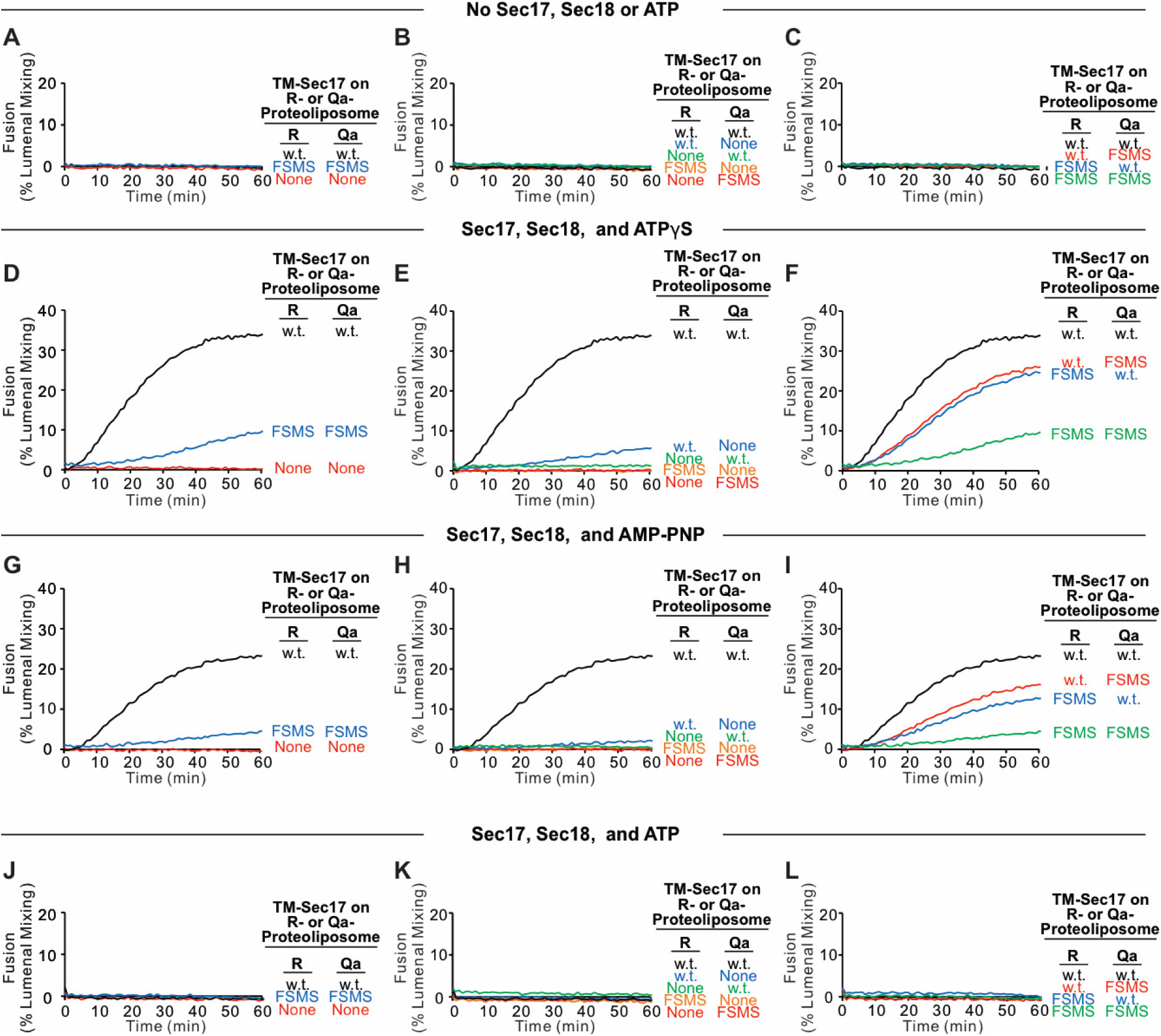
Fusion with membrane-anchored Sec17. Fusion was assayed between proteoliposomes bearing R, Ypt7, and either no Sec17, TM-anchored Sec17, or TM-anchored F22SM23S Sec17 and proteoliposomes with Qa, Ypt7, and either no Sec17, TM-anchored Sec17, or TM-anchored F22SM23S Sec17. Ypt7 was present at 1:8,000 molar ratio to lipid, and SNAREs and TM-Sec17 at 1:16,000 molar ratio to lipid. Fusion reactions had the indicated proteoliposomes, 50nM HOPS, 2μM each of sQb3Δ and Qc3Δ, and either **A-C** Sec18 buffer, **D-F** 1mM ATPγS and Sec18, **G-I** 1mM AMP-PNP and Sec18, or **J-L** 1mM ATP and Sec18.

### Selective inhibitors reveal successive fusion intermediates

Ypt7/R- and Ypt7/Qa-proteoliposomes were co-incubated in the presence of HOPS and other fusion proteins or inhibitors, allowing resolution of an early, HOPS-dependent reaction stage from late, Sec18-dependent fusion. Incubations were in sets of 3, with either of 3 antibody preparations. When HOPS, Ypt7/R- and Ypt7/Qa-proteoliposomes were mixed with sQb3Δ, Qc3Δ, Sec17, Sec18, and ATPγS plus control IgG or specific antibody from the start of the incubation, fusion was blocked by affinity-purified antibody to either Sec18 (Haas and Wickner, 1996) or to Vps33 (Seals et al., 2000), the SM subunit of HOPS (Figure 9A, Set 1). When the proteoliposomes were incubated for 29 minutes with HOPS, sQb3Δ, and Qc3Δ prior to antibody addition, followed by Sec17, Sec18, and ATPγS one minute later, the restored fusion remained sensitive to each affinity-purified antibody (Set 2). If however the initial 29 minute incubation also bore Qc3Δ, alone (Set 4) or in combination with other proteins (Sets 5-8), the restored fusion had acquired resistance to antibody to Vps33 prior to addition of the missing fusion proteins. Incubation with sQb3Δ did not confer resistance to anti-Vps33 (Set 3). Sec17 and Sec18 are not needed to form a reaction intermediate which is resistant to antibody to Vps33 (Set 4). In contrast, when fusion reactions were blocked by the single omission of either sQbΔ (Set 5) or Qc3Δ (Set 3), any intermediates which formed were not resistant to antibody to Sec18. A very partial resistance was seen when all 4 SNAREs were present during the initial 29 minute incubation (Sets 7, 8). The specificity of inhibition by antibody to Sec18 was confirmed by its failure to inhibit Sec18-independent fusion with full-length sQb and Qc (Figure 9B, C) but there was complete inhibition of Sec18-dependent fusion reactions with sQb3Δ and Qc3Δ (Figure 9D and E, red). Sec18 thus acts late in the fusion pathway, as incubations of the proteoliposomes with HOPS and Qc3Δ, which confers fusion-resistance to antibody to Vps33 (Figure 9, Set 4), remain sensitive to antibody to Sec18.

**Figure 9.**
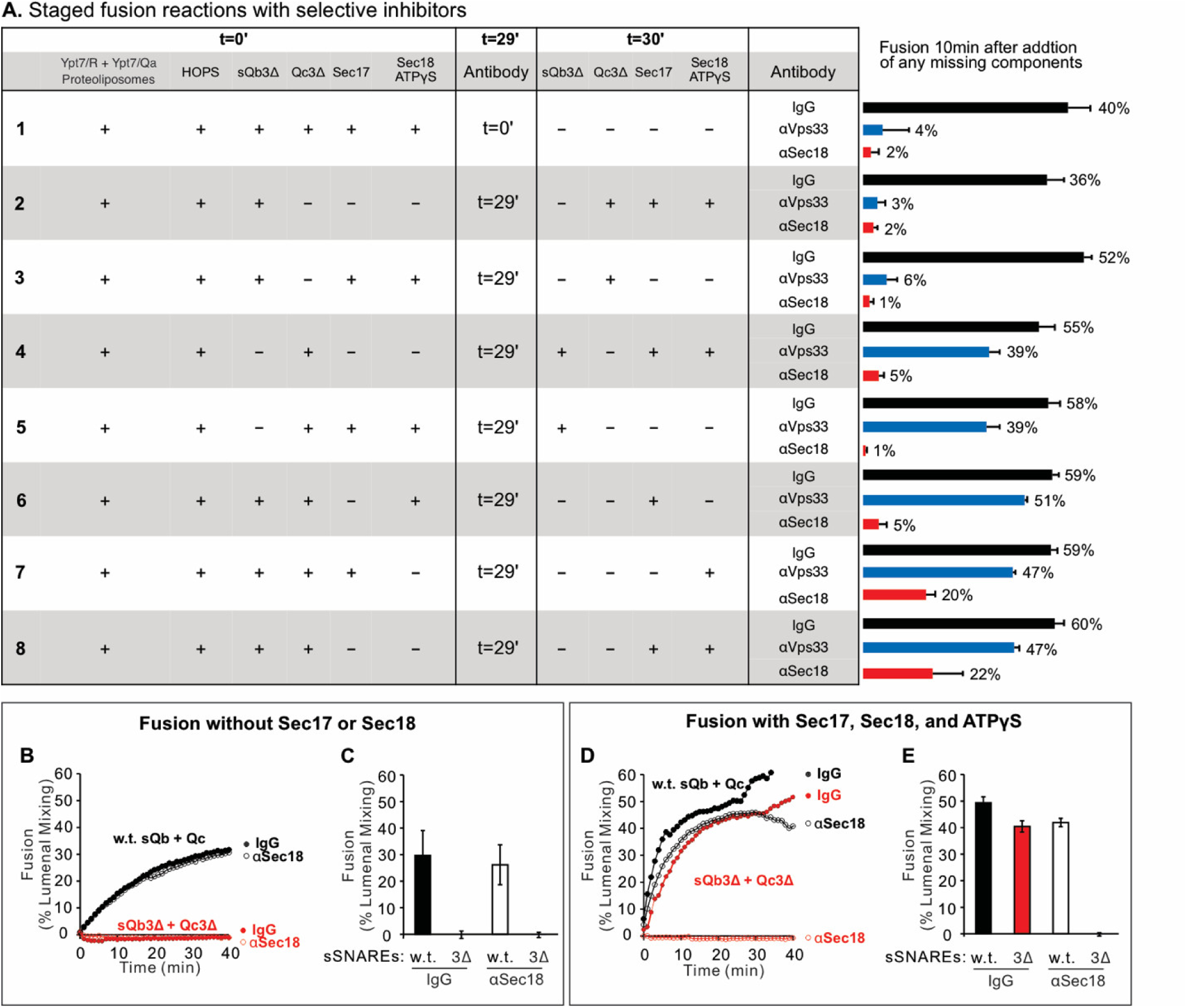
Stages of the fusion reaction, determined by the order of addition of fusion proteins and selective inhibitors. **A**. Proteoliposomes with Ypt7/R and proteoliposomes with Ypt7/Qa (1:8,000 Ypt7-TM:lipid molar ratio, 1:16,000 SNARE:lipid molar ratios) were mixed with 50 nM HOPS at t=0. At either t = 0 or at t = 30, indicated subsets of 2μM sQb3Δ, 2μM Qc3Δ, 400nM Sec17, 300nM Sec18, and 1mM ATPγS were added to the incubations. To block Sec18 or Vps33, affinity-purified antibody (1μg) to each was added at t=0 (sample 1) or at t=29 (samples 2-8). As a control, 1μg IgG was added to separate samples as indicated. The average and standard deviations of fusion 10 min after addition of any missing components is shown from three independent experiments. **B-E**. Sec18 antibody inhibition experiments were performed with Ypt7/R and Ypt7/Qa proteoliposomes with 50 nM HOPS, 2μM sQb and 2μM Qc (black lines) or 2μM sQb3Δ and 2μM Qc3Δ (red lines), 1μg IgG (filled circles) or αSec18 (open circles), and without (B and C) or with (D and E) 400nM Sec17, 300nM Sec18 and 1mM ATPγS as indicated. Kinetics shown are representative of 3 experiments.

### Fusion despite triply-crippled SNARE zippering

Since SNARE zippering is driven by the sequestration of apolar residues into the interior of the 4-SNARE bundle, we examined the effect of converting the apolar residues of the Qa SNARE domain +4 to +8 layers to Ala, to Ser, or to Gly. Fusion between Ypt7/R- and Ypt7/Qa-proteoliposomes in the presence of HOPS, sQb, and Qc but without Sec17 or Sec18 (Figure 10A, black curve), was diminished by replacing each of the +4 to +8 layer apolar residues of Qa with Ala (Figure 10A, green curve) and was abolished when they were replaced by Ser (blue curve) or Gly (red curve). The persistence of some fusion with the Ala substitutions may reflect that two of the residues were already Ala, and that Ala has the greatest propensity among the amino acids to form α-helices, Gly the least, and Ser is inbetween (Pace and Scholtz, 1998). When these same incubations were performed with Sec17, Sec18, and ATPγS, rapid and comparable fusion was seen in each case (Figure 10B). When hydrolyzable ATP was used instead of ATPγS, there was little effect on the fusion kinetics with wild-type SNARE domain sequences (Figure 10C vs 10B, black lines). In contrast, hydrolyzable ATP led to fusion inhibition when SNARE packing stability was reduced by substitution of apolar residues by Ala, Ser, or Gly (Figure 10C). Though the apolar residues are not required for fusion (Figure 10B), they apparently stabilize the SNAREs against ATP-driven proofreading disassembly (Figure 10C). To triply weaken the completion of zippering, the same proteoliposomes with wild-type Qa or the Qa +4 to +8 layer having small side-chain residues instead of apolar residues were incubated with HOPS, sQb3Δ and Qc3Δ, either without Sec17 or Sec18 (Figure 10D) or with Sec17, Sec18, and ATPγS (Figure 10E) or ATP (Figure 10F). Fusion was optimally supported by Sec17, Sec18, and ATPγS (Figure 10E). The independence of this fusion from energy derived by zippering is underscored by the similar fusion rates in all incubations with Sec17, Sec18, and ATPγS, whether with apolar or polar Qa +4 to +8 residues or with full-length or +4 to +8-truncated sQb and Qc (Figures 10B vs. E). With the Qb and Qc SNARE domains truncated, and the Qa lacking apolar inward-facing amino acyl side-chains, little or no energy could be gained from the completion of R and mutant-Qa zippering. Thus Sec17 acts in two ways, both enhancing zippering in the presence of HOPS and full-length SNARE domains (Figure 3) and acting with Sec18 to trigger fusion independent of energy from zippering (Figure 10).

**Figure 10.**
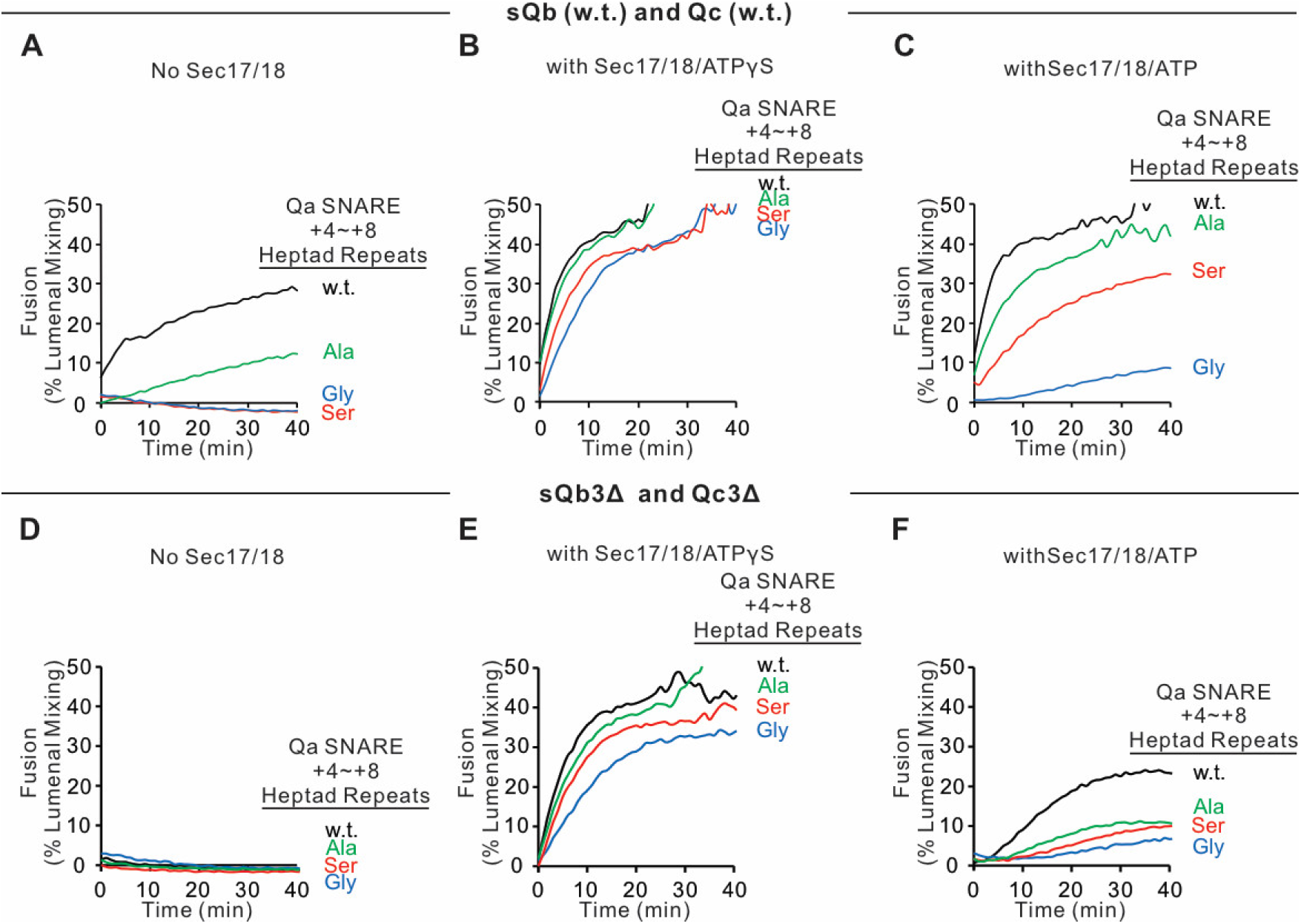
Sec17, Sec18, and ATPγS restore fusion to Ypt7/R and Ypt7/Qa proteoliposomes which were triply-blocked from completion of SNARE domain zippering by deletion of the +4 to +8 layers of the sQb and Qc SNAREs and by substitution of the apolar residues of the +4 to +8 layers of Qa, substituting Ala, Ser, or Gly for L238, M242, A245, L249, and A252. Fusion incubations (see Methods) had **A-C**. Ypt7/R- and Ypt7/Qa (w.t. (Black), Gly mutant (Blue), Ser mutant (Red) or Ala mutant (Green)) proteoliposomes (1:8,000 Ypt7-TM:lipid molar ratio, 1:16,000 SNARE:lipid molar ratios), 50nM HOPS, 2μM sQb (w.t.) and 2μM Qc (w.t.) (A - C) or **D-F** 2μM Qc3Δ and 2μM sQb3Δ. Sec17 and Sec18 buffers (A, D) or 600nM Sec17, 300nM Sec18, and 1mM ATPγS (B, E) or ATP (C, F) were also present. Kinetics shown are representative of 3 experiments.

## Discussion

The catalytic roles of fusion proteins have been gleaned from functional reconstitution studies. These studies initially showed that the zippering of concentrated SNAREs can drive slow fusion (Weber et al., 1998; Fukuda et al., 2000). As proteoliposome SNARE levels were lowered towards physiological levels, reconstituted vacuolar and neuronal fusion reactions required additional factors (Zick and Wickner, 2016; Stepien and Rizo, 2021). In addition to SNAREs, reconstituted neuronal fusion requires Munc18, Munc13, calcium, NSF, and SNAP (Ma et al., 2013; Lai et al., 2017) while reconstituted vacuole fusion needs HOPS (Stroupe et al., 2006), the Rab Ypt7 (Stroupe et al., 2009; Ohya et al., 2009), and specific lipid head-group composition and fatty acyl fluidity (Zick and Wickner, 2016). HOPS and other SM proteins catalyze SNARE assembly (Baker et al., 2015; Orr et al., 2017; Jiao et al., 2018), regulated by an activated Rab (Zick and Wickner, 2016; Torng and Wickner, 2020), and may confer resistance to Sec17/αSNAP and Sec18/NSF - mediated *trans*-SNARE disassembly (Xu et al., 2010; Jun and Wickner, 2019). Sec17/αSNAP and Sec18/NSF don’t simply regulate fusion, but actually stimulate fusion with HOPS and wild-type SNAREs (Mima et al., 2008; Song et al., 2017).

Fusion can be supported by either complete 4-SNARE zippering without Sec 17 or Sec18, or by Sec17 and Sec18 association with partially zippered SNAREs, but the most rapid fusion requires both (Song et al., 2017). While Sec17 and Sec18 are known to bypass the fusion blockade by Qc C-terminal truncation alone (Schwartz and Merz, 2009; Schwartz et al., 2017), the generality of this bypass with respect to any one Q-SNARE or even two Q-SNAREs (Figures 5,6) shows that it is not specific to Qc alone. Moreover, when zippering with full-length SNARE domains is weakened by the substitution of small amino acyl residues for large apolar residues in the +4 to +8 layers of Qa, Sec17 and Sec18 will also restore fusion (Figure 10 A, B). Strikingly, Sec17 and Sec18 drive fusion despite a triple blockade to complete zippering, namely the absence of two C-terminal Q-SNARE domains and the lack of apolarity in the third (Figure 10 C, D). In the NSF/αSNAP/SNARE complex, the 4-SNAREs wrap around each other in a left-handed super-helix, and the Sec17’s wrap around them in a right-handed fashion, yet they form a specific structure (Zhao et al., 2015; White et al., 2018). It seems unlikely that residues from one or more Sec17 could substitute for the missing residues when 2 heptads are removed from the C-termini of one or even two Q-SNAREs and the bulky apolar residues are removed from the third.

From the earliest reconstitutions of HOPS-dependent fusion (Mima et al., 2008) and in subsequent studies (e.g. Zick et al., 2015), Sec17 and Sec18 gave strong stimulation. In contrast, Sec17 and Sec18 inhibit SNARE-only fusion, or fusion with nonphysiological tethers (Mima et al., 2008; Zick and Wickner, 2014; Schwartz et al., 2017; Song and Wickner, 2019). HOPS not only binds each SNARE, but has direct affinity for Sec17 (Figure 4) and is necessary for Sec17 to enhance zippering per se (Figure 3C). This binding may underlie the HOPS:Sec17 synergy for fusion, but further studies are needed to test this idea.

Earlier studies and our current work suggest a working model of vacuole membrane fusion, encompassing findings here and elsewhere (Figure 11). HOPS exploits the affinity of its Vps39 and Vps41 subunits for the Rab Ypt7 (Brett et al., 2008) on each fusion partner membrane (Figure 11A) to mediate [step1] tethering (Hickey and Wickner, 2010). Tethered membranes (Figure 11B) are a prerequisite for SNARE assembly in an active conformation, likely a common N to C SNARE domain orientation (Song and Wickner, 2019). HOPS has direct affinity for each of the 4 vacuolar SNAREs (Song et al., 2020) and is allosterically activated by vacuolar lipids and Ypt7:GTP (Torng and Wickner, 2020) as a catalyst of SNARE assembly [step2]. SNAREs begin to zipper (Figure 11C) from the N towards the C-terminal end of their SNARE domain. When any one Q-SNARE is omitted, fusion intermediates assemble which undergo very rapid fusion when the missing Q-SNARE is supplied (Song et al., 2020), suggesting that fusion without Sec17/Sec18 is rate-limited by the completion of SNARE zippering and/or the spontaneous release of bulky HOPS. Sec17 has direct affinity for SNAREs (Söllner et al., 1993), HOPS (Figure 4), lipid (Clary et al., 1990; Zick et al., 2015), and Sec18 (Söllner et al., 1993). Sec17 displaces HOPS from the SNAREs [Figure 11, step 3], as shown by earlier studies: 1. Vacuolar SNAREs are found in complex with Sec17 or with HOPS, but not with both, and Sec17 can displace HOPS from SNAREs (Collins et al, 2005). 2. When vacuolar HOPS-dependent reconstituted fusion is arrested by Qc3Δ, HOPS is bound to the incompletely-zippered SNARE complex until it is displaced by Sec17 (Schwartz et al., 2017). 3. *trans*-SNARE complexes which assemble between isolated vacuoles are largely associated with Sec17 (Xu et al., 2010). The Sec17 associatiion with partially-zippered *trans*-SNARE complex (Figure 11D) promotes compete zippering (Figure 3) and will bind Sec18 [Step 4] through the Sec18 affinities for both SNAREs (Zick et al., 2015) and for Sec17. Sec18 regulates in some unknown fashion the Sec17/αSNAP assembly into an oligomeric cage surrounding the SNARE complex intermediate (Zhao et al., 2015), a *trans*-anchored Sec18/Sec17/SNARE pre-fusion complex (Figure 11E). Some ATP hydrolysis-dependent disassembly can occur, which may represent proofreading of incorrect and unstable *trans*-SNARE complexes. Sec17 oligomerization may also be stabilized or guided by the insertion of its N-terminal loop into the membranes. While SNAREs can slowly complete zippering and support fusion without Sec17 or Sec18 (Mima et al., 2008; Song et al., 2015), and slow fusion can occur with Sec17 and Sec18 when sQb and Qc lack the C-terminal portion of their zippering domain (Figure 6A), optimal fusion requires 4 complete SNARE domains, Sec17/αSNAP, and Sec18/NSF. The energy for fusion [step 5] derives from multiple sources: the completion of SNARE zippering, the binding energies which create the Sec17 cage for entropic confinement of the SNAREs, and from the energy of bilayer distortion through Sec17 apolar loop insertion. *Cis*-SNARE complexes (Figure 11F) formed by fusion (Söllner et al., 1993; Mayer et al., 1996) are disassembled by Sec17, Sec18, and ATP [step 6], freeing SNAREs from each other (Figure 11A) for later assembly in *trans*. Each of these components- the four SNAREs, the Rab Ypt7, HOPS, Sec17, and Sec18-are part of this ordered pathway of associations and disassembly, constituting a holoenzyme for vacuole membrane fusion.

**Figure 11.**
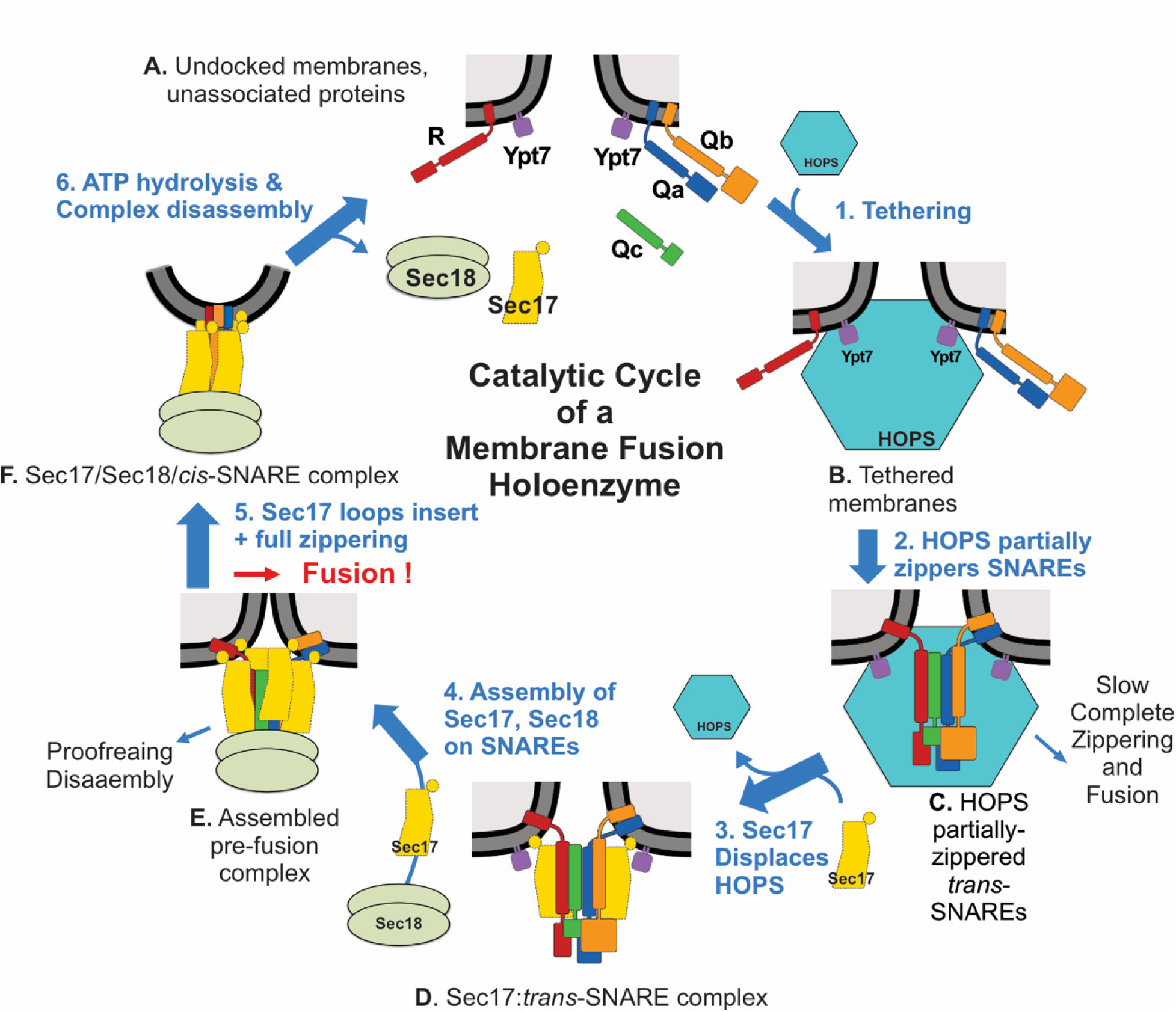
Current working model. Catalyzed tethering and SNARE assembly (A-C) is followed by HOPS displacement by Sec17 (C-D) and further assembly of Sec17 and Sec18 on the partially-zippered SNARE complex, promoting completion of zippering and apolar Sec17 loop insertion (D-F), both of which promote fusion. See text for discussion.

Our current studies reveal a general capacity of Sec17/αSNAP and Sec18/NSF to support fusion, even when little or no energy would be derived from completion of zippering. Zippering itself is enhanced by Sec17. An “entropic cage” (Chakraborty et al., 2010) formed by Sec17 molecules may favor SNARE zippering or, where zippering is blocked, allow the remaining SNARE domains to remain in positions and conformations which approximate zippering. The apolar N-domains of the four Sec17s are clearly essential for fusion (Figures 6, 7), whether through positioning the Sec17s, facilitating their assembly, or directly inserting as a “membrane wedge” and thereby contributing to bilayer disruption at the fusion site. The continued need for this apolar loop when Sec17 is integrally membrane-anchored (Figure 7) shows that it does not simply contribute to Sec17 membrane association. Elements of “entropic cage” and “membrane wedge” action are not mutually exclusive. Further work will be needed to determine the relative energies of Sec17 and Sec18 binding to the assembling 4-SNARE *trans*-complex, each of four Sec17s inserting its apolar loops into the membrane, Sec17 forming lateral associations with other Sec17 molecules in the cage around the SNAREs, and complete SNARE zippering, as each of these helps to achieve the bilayer distortions of fusion.

## Materials and Methods

PI3P was from Echelon (Salt Lake City, Utah), ergosterol from Sigma (St. Louis, MO), fluorescent lipids from Thermo Fisher (Waltham, MA), and other lipids were from Avanti (Alabaster, AL). Biobeads SM2 were from BioRad, Cy5-Streptavidin from SeraCare (Milford, MA), biotinylated phycoerythrin from Invitrogen (Eugene, OR) and underivatized streptavidin from Thermo Fisher. Spectrapor 6 dialysis tubing (7.5mm dia, 25 kDa cutoff) was from Spectrum Labs (Las Vegas, NV). Octyl-β-D-glucopyranoside was purchased from Anatrace (Maumee, OH).

## Mutant constructions

Sec17 with 6 acidic amino acids mutated to neutral residues, GST-Sec17 (E34S, E35S, D38S, E73A, D43A, E75A), was generated by PCR with Phusion high-fidelity DNA polymerase (NEB). The DNA fragment was cloned into BamHI and SalI digested pGST parallel1 vector (Sheffield et al.,1999) with an In-Fusion kit (Takara Bio USA, Mountain View, CA). Using inverse PCR, pParallel1-GST-Sec17 mutant (E34S, E35S and D38S) was amplified with Phusion high-fidelity DNA polymerase (NEB) from a GST-Sec17 construct. The amplified linear DNA was re-circularized with an In-fusion kit (Takara Bio USA, Mountain View, CA). To generate Sec17 acidic to neutral mutants, pParallel1-GST-Sec17 mutant (E34S, E35S, and D38S) was amplified with a Sec17 E73A, D43A, and E75A mutant primer set using Phusion high-fidelity DNA polymerase (NEB) and re-circularized with an In-fusion kit (Takara Bio USA, Mountain View, CA) For Sec17-E34S,E35S, D38S:

F TCGTCGGCTGCTTCTCTTTGTGTCCAAGCAGCCAC

R AGAAGCAGCCGACGAAAACTTGTATGAATCAGAAC

For Sec17 E73A, D43A, E75A:

F GGTAATGCAGCCGCAGCAGGAAATACCTACGTAGA

R TGCGGCTGCATTACCAGCCTTTTTCTGATAGTCAG

sQa3Δ, sQb3Δ. GST-sQa3Δ with amino acyl residues 1–235 and GST-sQb3Δ with amino acyl residues 1–160 were generated by PCR with Phusion high-fidelity DNA polymerase (NEB). DNA fragments were cloned into BamHI and SalI digested pGST parallel1 vector (Sheffield et al.,1999) with an In-Fusion kit (Takara Bio USA, Mountain View, CA).

For GST-sQa3Δ,

F: AGGGCGCCATGGATCCGATGTCCTTTTTCGACATCGA

R: AGTTGAGCTCGTCGACTAGATATTCTCGTCTATGGTGG

For GST-sQb3Δ,

F: AGGGCGCCATGGATCCGATGAGTTCCCTATTAATA

R: AGTTGAGCTCGTCGACTACAAGGTCTGTCTTGCATTTT

To generate Qa with L238, M242, A245, L249, and A252 changed to Ala, Ser or Gly, the pParallel1 GST vector with Qa lacking residues 228-257 was generated by inverse PCR with pParallel1 GST-Qa (Mima et al, 2008) using Phusion high-fidelity DNA polymerase (NEB). The DNA duplex with mutations (Gly, Ser or Ala) of the +4 to +8 heptad repeats was cloned into the amplified vector bearing Qa1-227 with an In-fusion kit (Takara Bio USA, Mountain View, CA).

For inverse PCR of pParallel1 GST and Qa 1-227

F: GACCAGCATCAGAGGGACCG

R: GTCTATGGTGGTTACTTGTT

For the Gly mutant of Qa

Sequence1:

GTAACCACCATAGACGAGAATATCTCGCATGGCCATGATAACGGCCAGAATGGCAACAAACAAGGCACCAGAGGCGACCAGCATCAGAGG

Sequence2:

CCTCTGATGCTGGTCGCCTCTGGTGCCTTGTTTGTTGCCATTCTGGCCGTTATCATGGCCATGCGAGATATTCTCGTCTATGGTGGTTAC

For the Ser mutant of Qa

Sequence1: GTAACCACCATAGACGAGAATATCTCGCATAGCCATGATAACAGCCAGAATAGCAACAAACAAAGCACCAGAAGCGACCAGCATCAGAGG

Sequence2:

CCTCTGATGCTGGTCGCTTCTGGTGCTTTGTTTGTTGCTATTCTGGCTGTTATCATGGCTATGCGAGATATTCTCGTCTATGGTGGTTAC

For the Ala mutant of Qa

Sequence1: GTAACCACCATAGACGAGAATATCTCGCATGCCCATGATAACGCCCAGAATGCCAACAAACAAGCCACCAGAGCCGACCAGCATCAGAGG

Sequence2:

CCTCTGATGCTGGTCGGCTCTGGTGGCTTGTTTGTTGGCATTCTGGGCGTTATCATGGGCATGCGAGATATTCTCGTCTATGGTGGTTAC

### Protein isolation

HOPS and prenyl-Ypt7 (Zick and Wickner, 2013), Ypt7-TM (Song et al., 2020), Sec17 (Schwartz and Merz, 2009), TM-anchored Sec17 and TM-anchored Sec17-F21SM22S (Song et al., 2017), Sec18 (Mayer et al., 1996), wild-type vacuolar SNAREs and his6-R (Mima et al., 2008; Schwartz and Merz, 2009; Zucchi and Zick, 2011; Izawa et al., 2012), sQb (Zick and Wickner, 2013) and Qc3Δ (Schwartz and Merz, 2009) were purified as described.

sQa, sQa3Δ, and sQb3Δ were purified by a modification of prior methods (Zick and Wickner, 2013; Song et al., 2020). pGST-Parallel1 with sQa, sQa3Δ, or sQb3Δ were transformed into Rosetta2 (DE3)-pLysS cells (EMD Millipore, Billerica, MA). Luria–Bertani (LB) broth (100ml) containing 100 μg/ml ampicillin and 34 μg/ml chloramphenicol was inoculated with a single colony. After overnight incubation with shaking at 37°C, 40ml portions of the preculture were added to two 6L flasks, each with 3L of LB medium, containing 100 μg/ml ampicillin and 34 μg/ml chloramphenicol and shaken (200rpm) at 37°C to an OD600 of 0.8. Isopropyl β-D-1-thiogalactopyranoside was added to 0.5 mM. After 3 h of continued shaking at 37°C, bacteria were harvested by centrifugation (5000rpm, 5 min, 4°C). Cell pellets were resuspended in 60 ml of 20 mM HEPES-NaOH (pH 7.4), 200 mM NaCl, 1 mM EDTA, 1mM DTT, 200mM phenylmethyl sulfonylfluoride, and 1 X protease inhibitor cocktail (Xu and Wickner, 1996). Resuspended cells were passed twice through a French press at 900 psi. The cell lysate was centrifuged (4°C, 1 h, 50,000 rpm, SW 60Ti rotor [Beckman Coulter, Brea, CA]). The supernatant was added to 20 ml of Glutathione Agarose 4B resin (Genesee Scientific, San Diego, CA) which had been equilibrated with wash buffer (20 mM HEPES-NaOH (pH 7.4), 200 mM NaCl, and 1 mM EDTA, 1mM DTT) and nutated at 4°C for 2 h. The suspended resin was poured into a 2.5-cm-diameter column, drained, and washed with 100ml wash buffer. The GST-tagged protein was eluted with 20 mM HEPES-NaOH (pH 7.4), 200 mM NaCl, 1 mM EDTA, 1mM DTT, 5% Glycerol, and 20mM glutathione, and the GST tag removed by TEV protease.

### Proteoliposome fusion

Proteoliposomes were prepared by detergent dialysis from β-octylglucoside mixed micelles for fusion assays (Song et al., 2017; Song et al., 2019) and SNARE assembly assays (Torng et al., 2020) as described. Briefly, for fusion assay, proteoliposomes (1 mM lipid) were prepared with membrane-bound Ypt7 and R at 1:8000 and 1:16,000 molar ratios to lipid, respectively, and with lumenal biotin-phycoerythrin. Proteoliposomes were also prepared with membrane-bound Ypt7 and the indicated Q-SNAREs at 1:8000 and 1:16,000 molar ratios to lipid, respectively, plus lumenal Cy5-streptavidin. These were incubated separately for 10min at 27°C with 1μM GTP and 1 mM EDTA, then MgCl2 was added to 2mM. After prewarming (27°C 10min) the separately GTP-exchanged proteoliposomes in a 384 well plate, fusion reactions were initiated by mixing 5ul of each proteoliposome preparation and supplementing with other fusion factors in volumes summing to 10 μl, continuing incubation at 27°C in a Spectramax fluorescent plate reader. Fusion incubations (20 ul) in RB150 (20mM HEPES/NaOH, pH 7.4, 150 mM NaCl, 10% glycerol) had proteoliposomes (0.5mM total lipid concentration), 50nM HOPS, the indicated concentrations of sSNAREs, 400 or 600nM Sec17, 300nM Sec18, 1mM ATP or its analogs, and 3mM MgCl2, as modified in each Figure legend.

### SNARE assembly assay

Assays were performed as described previously (Torng et al., 2020) with one addendum. In brief, reactions (20 µL) were at 27 °C in a SpectraMax Gemini XPS (Molecular Devices) plate reader. Standard reactions include HOPS (160 nM), proteoliposomes [0.5 mM lipid, with SNARE and Ypt7 at a molar ratios of either 1:2000 or 1:4000 for Ypt7/R-proteoliposomes and Ypt7/RQa-proteoliposomes, respectively], and fluorescently-labeled Qb and Qc (1 µM), and sQa (1 µM) as necessary. These were incubated for 60 min, then Sec17 was added to 500nM. Three fluorescence channels were read simultaneously at intervals of 30 s: the donor channel Oregon Green 488 from Qc (excitation [ex]: 497 nm; emission [em]: 527 nm; cutoff [c/o]: 515 nm), the acceptor channel Alexa Fluor 568 from Qb (ex: 568 nm; em: 605 nm; c/o: 590 nm), and the FRET channel (ex: 490 nm; em: 615 nm; c/o: 590 nm). For each timepoint, the bleed through-corrected FRET signal was obtained by subtracting the background signals coming from the donor and acceptor channels from the signal in the FRET channel as detailed in Torng et al. (2020). This was further corrected by dividing by the geometric mean of the donor and acceptor signals. The final corrected signal, reported as “Average FRET efficiency,” is a combined measure of the proportion of fluorescent SNAREs undergoing FRET and their average FRET efficiency.

### Molecular models

MODELLER (Webb & Sali, 2016) was used to create individual homology models of the vacuolar SNARE complex (Nyv1, Vam3, Vti1, Vam7), and of Sec18 starting from the coordinates of synaptobrevin-2, SNAP-25, syntaxin-1A, and NSF in the cryo-EM structure of the neuronal NSF/αSNAP/SNARE complex (PDB ID 3J96) (Zhao et al., 2015). For Sec18, the linker between the N and D1 domains was deleted from the generated homology model since there was no information about these linkers in this structure (PDB ID 3J96) of the neuronal 20S complex.

The MODELLER protocol consisted of an alignment step (python script file align.py) and a modelling step (python script file modeler-input.py). The script files are shown here for synaptobrevin (nyv1):

#### align.py

from modeller import *

env = environ()

aln = alignment(env)

mdl = model(env, file=’sb.pdb’, model_segment=(’FIRST:K’,’LAST:K’))

aln.append_model(mdl, align_codes=’sbK’, atom_files=’sb.pdb’)

aln.append(file=’target_sequence.pir’, align_codes=’nyv1’)

aln.salign(local_alignment=True, rr_file=’${LIB}/blosum62.sim.mat’,

gap_penalties_1d=(-600, -600),

output=’’,

align_block=15, # no. of seqs. in first MSA

align_what=’PROFILE’,

alignment_type=’PAIRWISE’,

comparison_type=’PSSM’, # or ’MAT’ (Caution: Method NOT benchmarked

\# for ’MAT’)

similarity_flag=True, # The score matrix is not rescaled

substitution=True, # The BLOSUM62 substitution values are

\# multiplied to the corr. coef.

#write_weights=True,

#output_weights_file=’test.mtx’, # optional, to write weight matrix

smooth_prof_weight=10.0) # For mixing data with priors

aln.edit(edit_align_codes=’nyv1’, base_align_codes=’rest’,min_base_entries=1, overhang=0)

aln.write(file=’nyv1.ali’, alignment_format=’PIR’)

aln.write(file=’nyv1.pap’, alignment_format=’PAP’)

#### modeler-input.py

from modeller import *

from modeller.automodel import *

env = environ()

a = automodel(env, alnfile=’nyv1.ali’,

knowns=’sbK’, sequence=’nyv1’,

assess_methods=(assess.DOPE, assess.GA341))

a.very_fast()

a.starting_model = 1

a.ending_model = 1

a.make()

The homology models of Nyv1, Vam3, Vti1, Vam7 and Sec18, together with the crystal structure of Sec17 (PDB ID 1QQE) (Rice & Brunger, 1999) were fit into the Cryo-EM structure of the neuronal NSF/αSNAP/SNARE complex (PDB ID 3J96) (Zhao et al., 2015). The fit was performed by using the “align” feature of PyMol to individually superimpose the coordinates of the vacuolar proteins with the corresponding coordinates of the neuronal proteins in the structure of the neuronal NSF/αSNAP/SNARE complex.

## Acknowledgements

This work was supported by grants R35GM118037 (to W.W.) and R37MH63105 (to A.T.B.) from the NIH.

---

**Figure 2-figure supplement 1.**
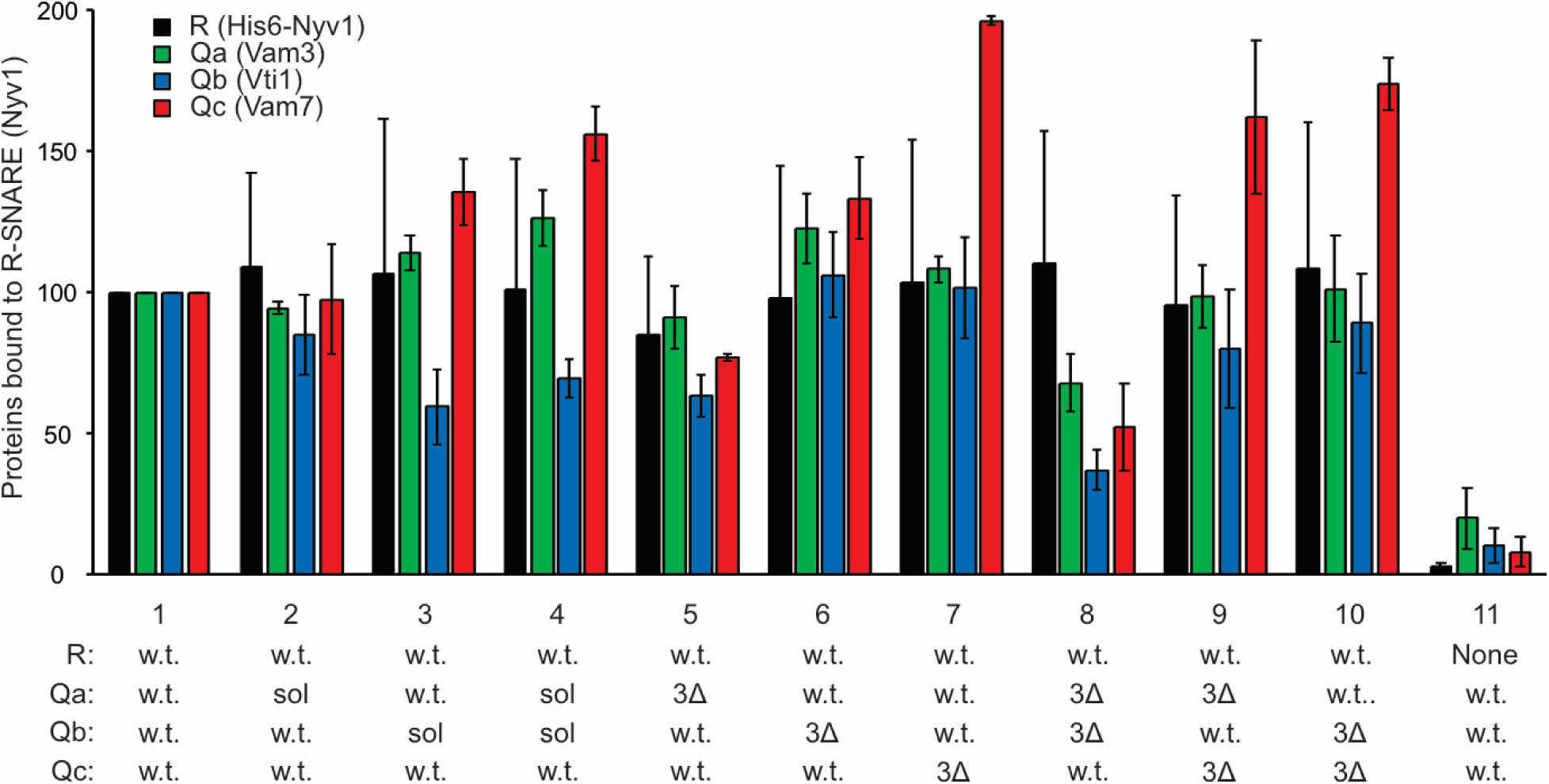
Stable associations of truncated SNAREs. Western blots of three identical experiments were scanned and analyzed with UN-SCAN-IT Software (Silk Scientific, Orem UT). Average pixel values were set at 100 for the all-wild-type positive control (lane 1). Standard deviations are shown.

**Figure 3-figure supplement 1.**
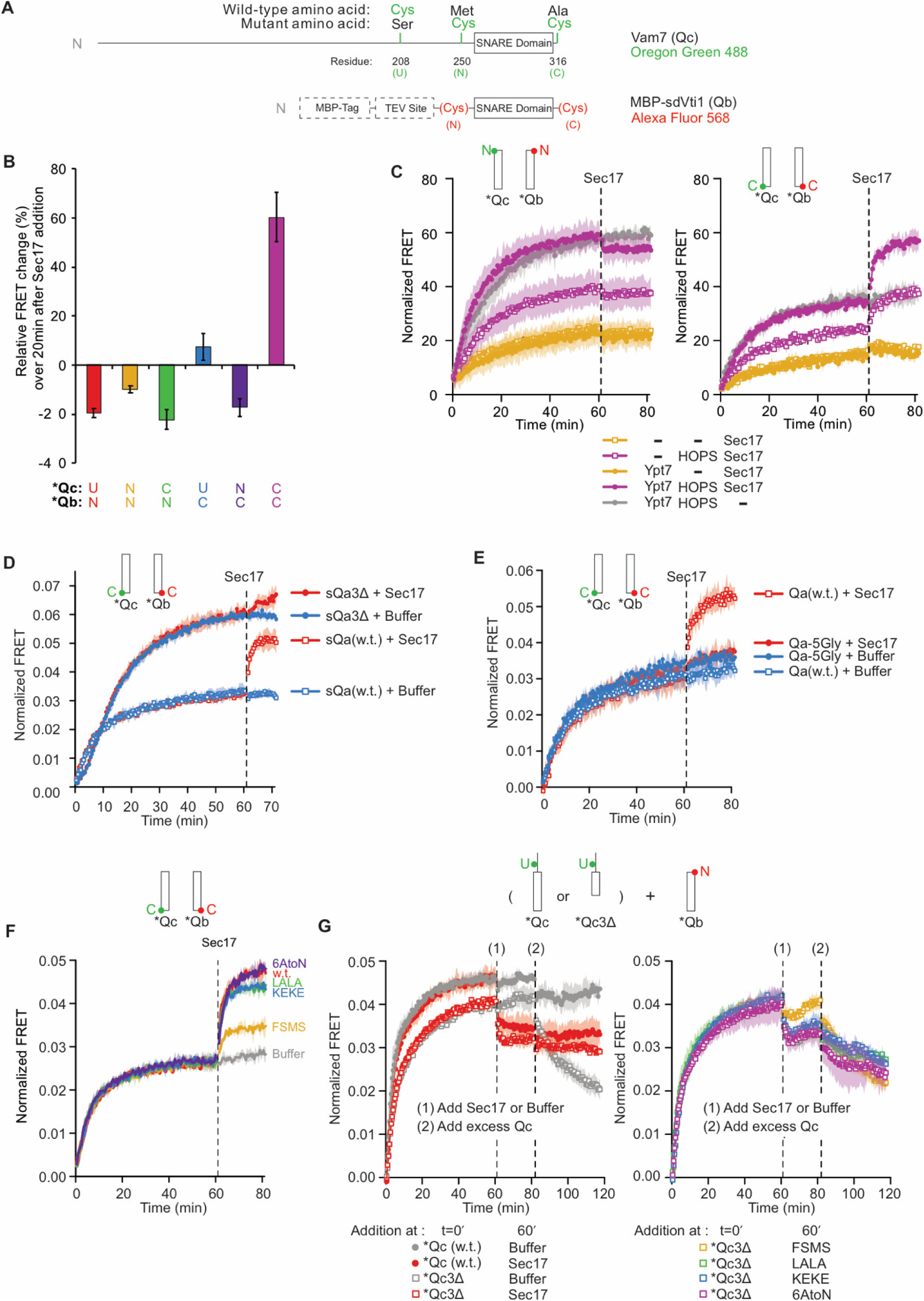
Statistics and representative kinetic data for Figure 3. **A**. Schematic of fluorescently labeled SNARE constructs used in this study. **B-G**. Supplementary data for panels B-G in Figure 3. Subfigures are presented with the same subfigure labels and matching colors. All kinetic data are averages of 3 trials, and the shaded regions behind each curve show the standard deviation per time point. **B**. The effect of Sec17 on SNARE average FRET efficiency as a function of fluorophore position. The mean and standard deviation of three trials is shown for each reaction. **C**. Representative kinetic data showing the effect of HOPS and Ypt7 on Sec17-induced zippering. Data for the N-terminally labeled *Qb/*Qc pair is shown on the left, and data for the C-terminally labeled pair is shown on the right. Reactions with HOPS are in purple, and reactions without HOPS are in orange. As a negative control, buffer (RB150) was added instead of Sec17 for the reaction in gray. **D**. Representative kinetic data showing that Sec17 has only minor effect on the average FRET efficiency of SNARE complexes bearing truncated Qa SNARE. **E**. Representative kinetic data showing that Sec17 does not affect the average FRET efficiency of SNARE complexes bearing Qa-SNARE with a mutated C-terminal half. **F**. Representative kinetic data for the ability of Sec17 or mutants to promote C-terminal zippering of the SNARE complex. **G**. Representative kinetic data showing that Sec17 stabilizes incompletely-zippered SNARE complexes. Sec17, mutants, or buffer was added at t = 60 min (1), and non-fluorescent Qc was added at t = 80 min (2). Reactions containing wildtype Qc are shown with filled circles, and reactions with Qc3Δ are shown with open squares. The effect of wildtype Sec17 (red) is contrasted with no Sec17 (gray) in the left-side graph, and the four Sec17 mutants used in this experiment are compared in the right-side graph.

**Figure 5-figure supplement 1.**
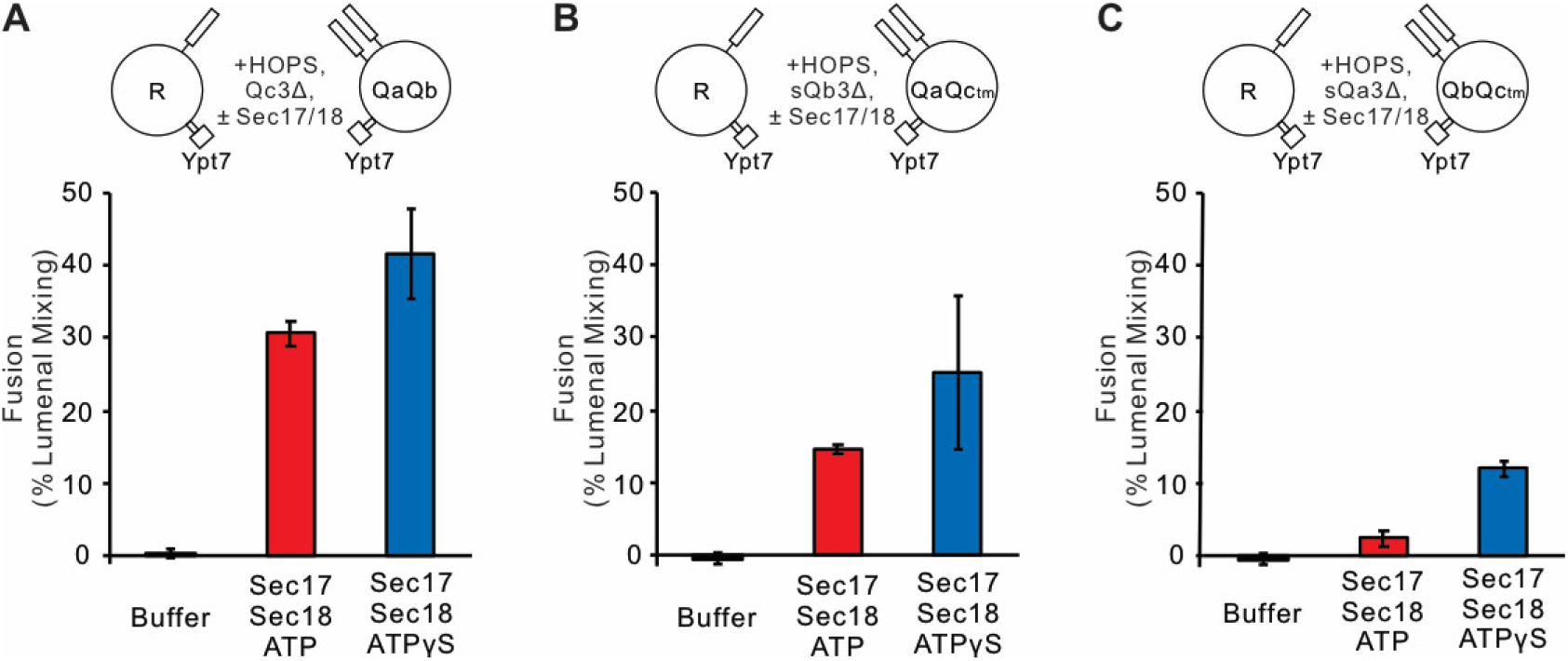
Statistics for Figure 5. Fusion blocked by deletion of the C-terminal 5 layers of any Q-SNARE domain is restored by Sec17 and Sec18. **A.** Ypt7/R- and Ypt7/QaQb, **B.** Ypt7/R- and Ypt7/QaQc-tm, and **C.** Ypt7/R- and Ypt7/QbQc-tm. The experiment in Figure 5 was repeated in triplicate; mean values and standard deviations for fusion after 30 minutes are shown.

**Figure 6-figure supplement 1.**
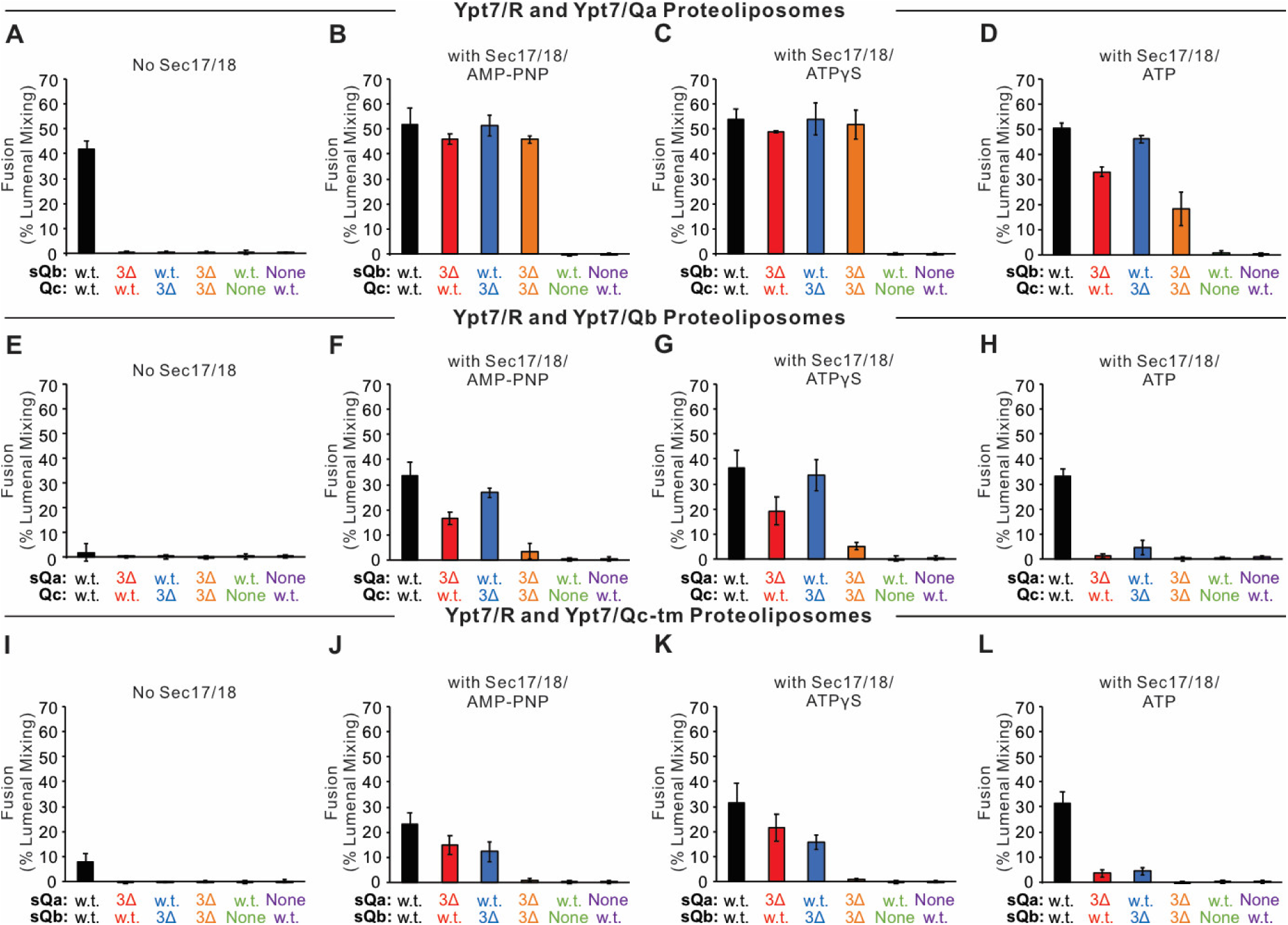
Statistics data for Figure 6. Fusion between Ypt7/R- and **A-D.** Ypt/Qa, **E-H.** Ypt7/Qb, or **I-L.** Ypt7/Qc-tm proteoliposomes. Fusion assays were described in Figure 6, repeated more than 3 times. Mean values and standard deviations for fusion after 20 minutes are shown.

**Figure 7-figure supplement 1.**
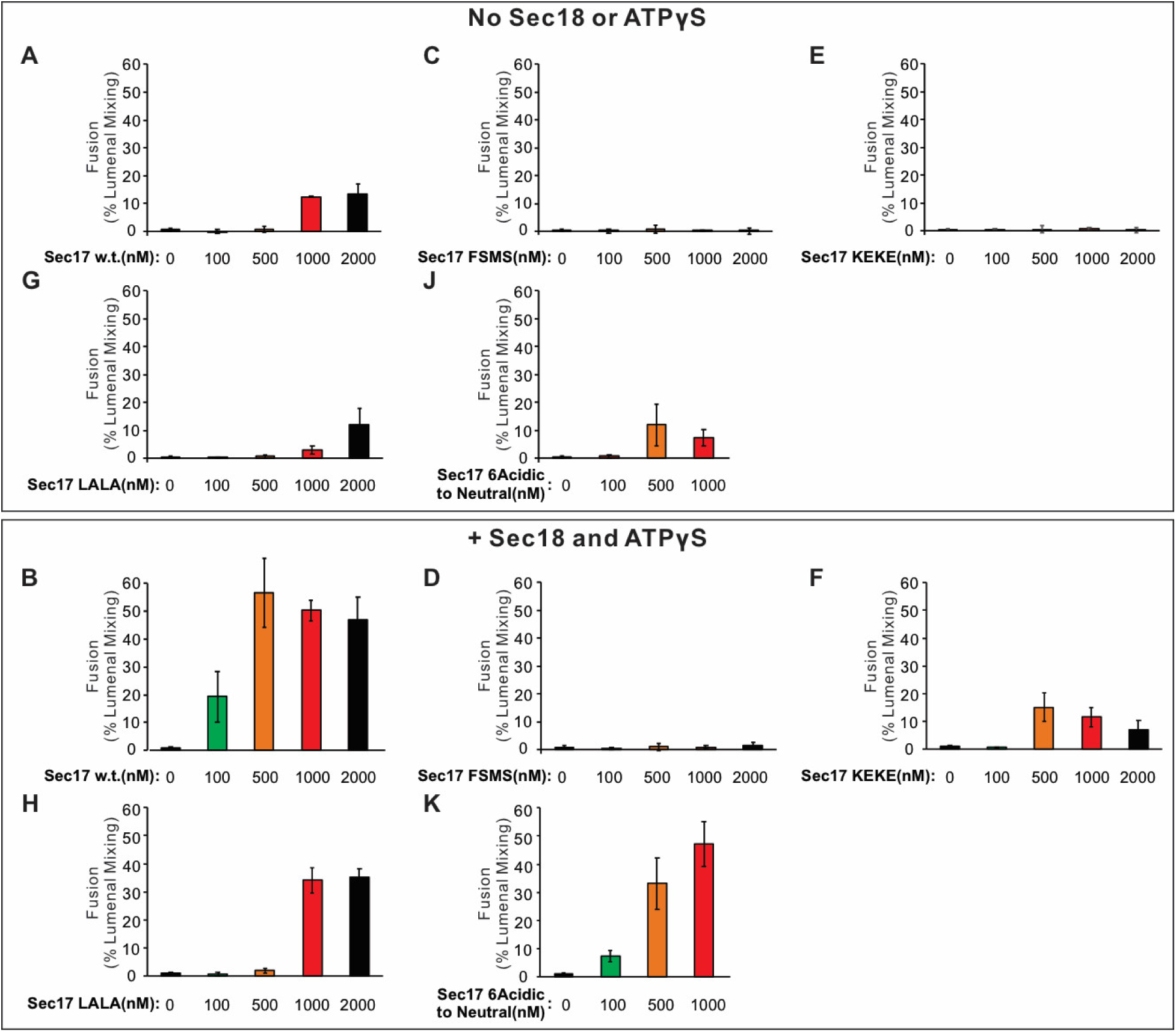
The experiment in Figure 7 was repeated in quadruplicate; mean values and standard deviations for fusion after 30 minutes are shown.

**Figure 8-figure supplement 1.**
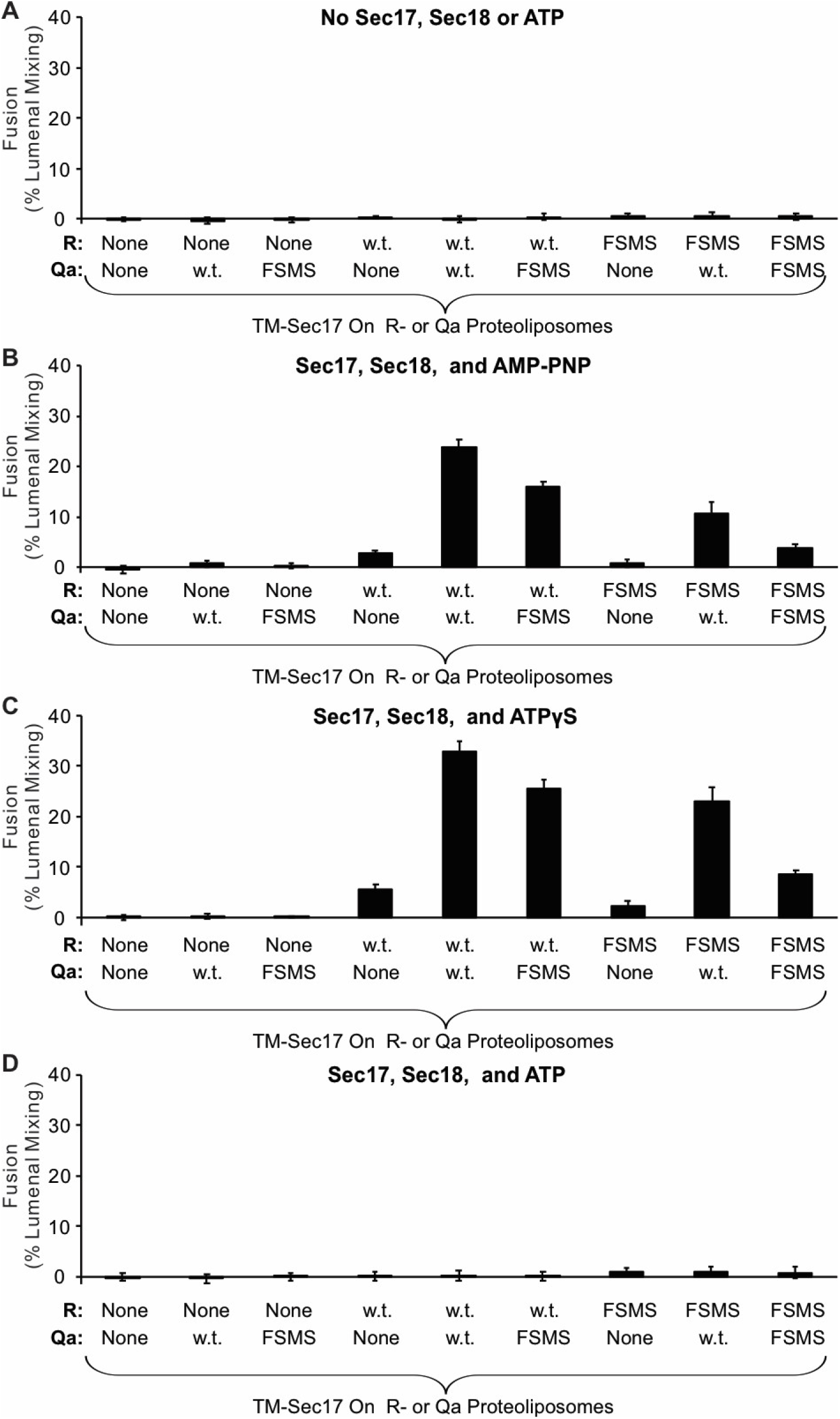
The experiment in Figure 8 was repeated in quadruplicate; mean values and standard deviations for fusion after 60 minutes are shown.

**Figure 9-figure supplement 1.**
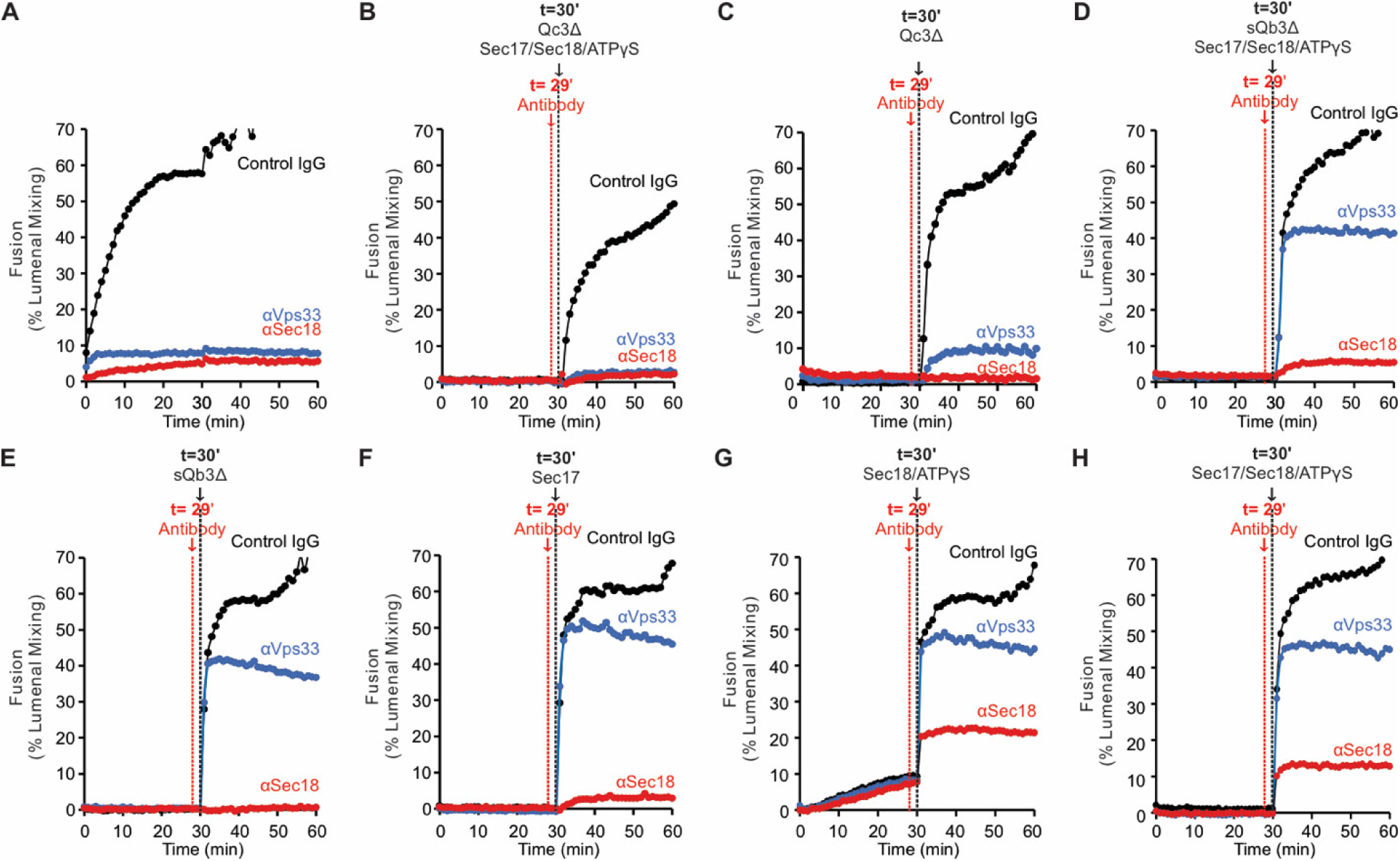
Detailed fusion kinetics for a representative experiment from Figure 9.

**Figure 10-figure supplement 1.**
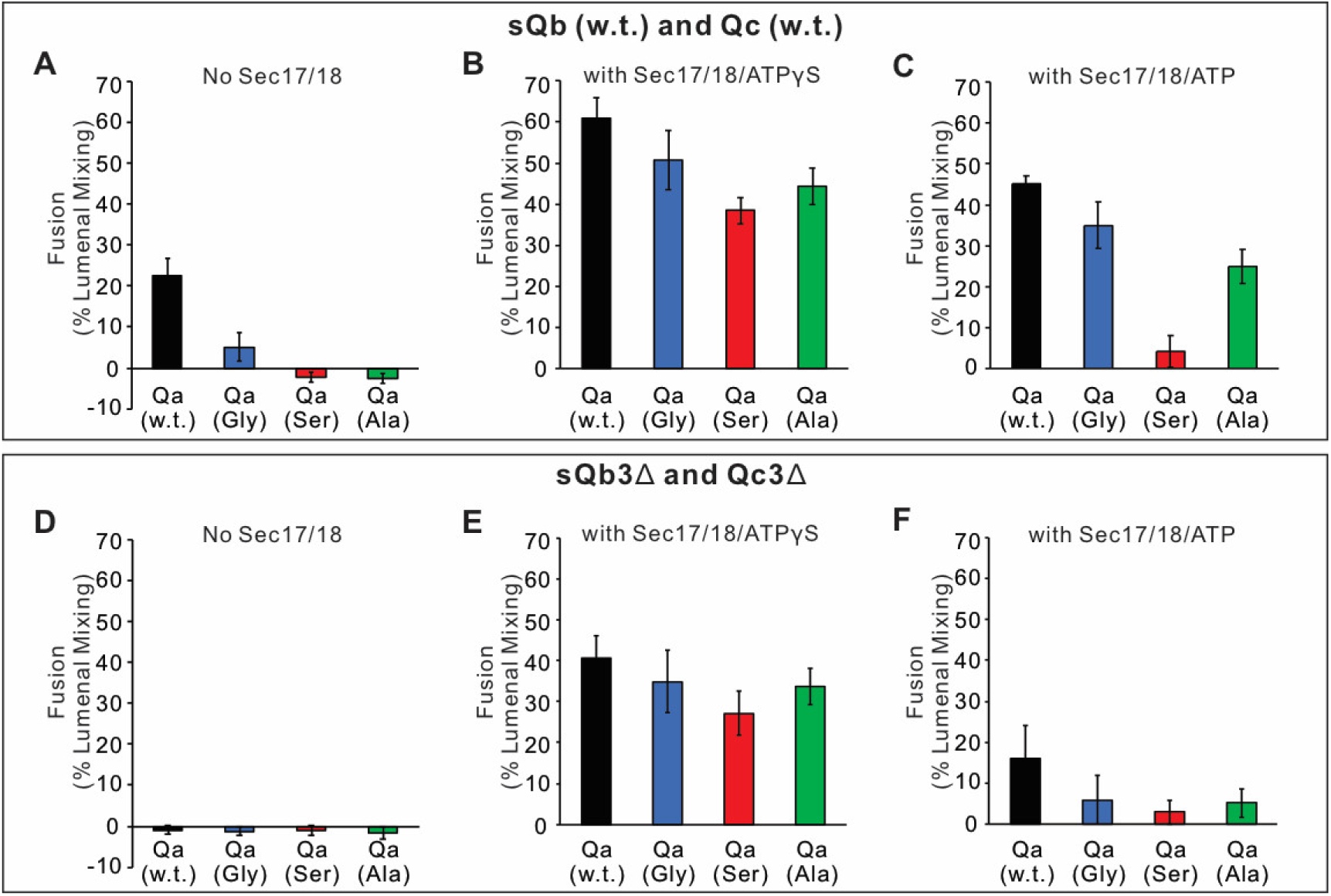
Fusion assays between Ypt7/R- and Ypt7/Qa with either Qa W.T. (Black), Qa Gly mutant (Blue), Qa Ser mutant (Red), or Qa Ala mutant (Green) were as described in Figure 10. **A-C.** Fusion with 2μM sQb and 2μM Qc. **D-F.** Fusion with 2μM sQb3Δ and 2μM Qc3Δ. Experiments were repeated in quadruplicate; mean values and standard deviations for fusion after 30 minutes are shown.

## References

1. Aeffner, S., Reusch, T., Weinhausen, B., and Salditt, T. (2012) Energetics of stalk intermediates in membrane fusion are controlled by lipid composition. Proc. Natl. Acad. Sci USA DOI 10.1073/pnas.1119442109

2. Baker, R.W., Jeffrey, P.D., Zick, M., Phillips, B.P, Wickner, W.T., and Hughson, F.M. (2015) A direct role for the Sec1/Munc18-family protein Vps33 as a template for SNARE assembly. Science 349, 1111–1114

3. Barnard, R.J., Morgan, A., and Burgoyne, R.D. (1997) Stimulation of NSF ATPase activity by alpha-SNAP is required for SNARE complex disassembly and exocytosis. J. Cell Biol. 139, 875–883.

4. Brett, C.L., Plemel, R.L., Lobinger, B.T., Vignali, M., Fields, S., and Merz, A.J. (2008) Efficient termination of vacuolar Rab GTPase signaling requires coordinated action by a GAP and a protein kinase. J. Cell Biol. 182, 1141–1151.

5. Chakraborty, K., Chatila, M., Sinha, J., Shi, Q., Poshner, B.C., Sikor, M., Jiang, G., Lamb, D.C., Hartl, F.U., and Hayer-Hartl, M. (2010) Cheronin-catalyzed rescue of kinetically trapped states in protein folding. Cell 142, 112–122.

6. Cheever, M.L., Sato, T.K., de Beer, T., Kutatelaade, T.G., Emr, S.D., and Overduin, M. (2001) Phox domain interaction with PtdIns(3)P targets the Vam7 t-SNARE to vacuole membranes. Nat. Cell Biol. 3, 613–618.

7. Choi, U.B., Zhao, M., White, K.I., Pfuetzner, R.A., Esquivies, L., Zhou, Q., Brunger, A.T. (2018) NSF-mediated disassembly of on- and off-pathway SNARE complexes and inhibition by complexin. DOI: 10.7554/eLife.36497.

8. Clary, D.O., Griff, I.C., and Rothman, J.E. (1990) SNAPs, a family of NSF attachment proteins involved in intracellular membrane fusion in animals and yeast. Cell 18, 709–721.

9. Collins, K.M., Thorngren, N.L., Fratti, R.A., and Wickner, W.T. (2005) Sec17p and HOPS, in distinct SNARE complexes, mediate SNARE complex disruption or assembly for fusion. EMBO J. 24, 1775–1786.

10. Fasshauer, D., Bruns, D., Shen, B., Jahn, R., and Brünger, A.T. (1997) A structural change occurs upon binding of syntaxin to SNAP-25. J. Biol. Chem. 272, 4582–4590.

11. Fasshauer, D., Sutton, R.B.B., Brunger, A.T.T., and Jahn, R. (1998) Conserved structural features of the synaptic fusion complex: SNARE proteins reclassified as Q-and R-SNAREs. Proc. Natl. Acad. Sci. USA 95, 15781–15786.

12. Fiebig, K.M., Rice, L.M., Pollock, E., and Brunger, A.T. (1999) Folding intermediates of SNARE complex assembly. Nat. Struct. Biol. 6, 117–123.

13. Fukuda, R., McNew, J.A., Weber, T., Parlati, F., Engel, T., Nickel, W., Rothman, J.E., and Söllner, T.H. (2000) Functional architecture of an intracellular membrane t-SNARE. Nature 407, 198–202

14. Gao, Y., Zorman, S., Gundersen, G., Xi, Z., Ma, L, Sirinakis, G, Rothman, J.E., and Zhang, Y. (2012) Single reconstituted neuronal SNARE complexes zipper in three distinct stages. Science 337, 1340–1343. DOI 10.1126/science.1224492

15. Haas, A. and Wickner, W. (1996) Homotypic vacuole fusion requires Sec17p (yeast α-SNAP) and Sec18p (yeast NSF). EMBO J. 15, 3296–3305.

16. Hazzard, J., Südhof, T.C., and Rizo, J. (1999) NMR analysis of the structure of synaptobrevin and of its interaction with syntaxin. J. Biomol. NMR 14, 203–207.

17. Hickey, C.M. and Wickner, W. (2010) HOPS initiates vacole docking by tethering membranes before trans-SNARE complex assembly. Mol. Biol. Cell 21, 2297–2305.

18. Izawa, R., Onoue, T., Furukawa, N., and Mima, J. (2012) Distinct contributions of vacuolar Qabc and R-SNARE proteins to membrane fusion specificity. J. Biol. Chem. 287, 3445–3453.

19. Jiao, J., He, M., Port, S.A., Baker, R.W., Xu, Y., Qu, H., Xiong, Y., Wang, Y., Jin, H., Eisemann, T.J., Hughson, F.M., and Zhang, Y. (2018) Munc18-1 catalyzes neuronal SNARE assembly by templating SNARE association. eLife DOI: https://doi.org/10.7554/eLife.41771.

20. Jun, Y. and Wickner, W. (2019) Sec17 (α-SNAP) and Sec18 (NSF) restrict membrane fusion to R-SNAREs, Q-SNAREs, and SM proteins from identical compartments. Proc. Natl. Acad. Sci. USA doi/10.1073/pnas.1913985116.

21. Lai, Y., Choi, U.B., Leitz, J., Rhee, H.J., Lee, C., Altas, B., Zhao, M., Pfuetzner, R.A., Wang, A.L., Brose, N., Rhee, J, and Brunger, A.T. (2017) Molecular mechanisms of synaptic vesicle priming by Munc13 and Munc18. Neuron 95, 591–607.

22. Ma, C., Li, W., Xu, Y., and Rizo, J. (2011) Munc13 mediates the transition from the closed syntaxin-Munc18 complex to the SNARE complex. Nat. Struct. Mol. Biol. 18, 542–549.

23. Ma, C., Su, L., Seven, A.B., Xu, Y., and Rizo, J. (2013) Reconstitution of the vital functions of Munc18 and Munc13 in neurotransmitter release. Science 239, 421–425.

24. Ma, L., Kang, Y., Jiao, J., Rebane, A.A., Cha, H.K., Xi, Z., Qu, H., and Zhang, Y. (2016) Alpha-SNAP enhances SNARE zippering by stabilizing the SNARE four-helical bundle. Cell Rep. 19, 531–539.

25. Marz K.E., Lauer, J.M., and Hanson, P.I. (2003) Defining the SNARE complex binding surface of alpha-SNAP: implications for SNARE complex disassembly. J. Biol. Chem. 278, 27000–27008.

26. Mayer, A., Wickner, W., Haas, A. (1996) Sec18p (NSF)-driven release of Sec17p (α-SNAP) can precede docking and fusion of yeast vacuoles. Cell 85, 83–94.

27. Mima, J., Hickey, C.M., Xu, H., Jun, Y., and Wickner, W. (2008) Reconstituted membrane fusion requires regulatory lipids, SNAREs, and synergistic SNARE chaperones. EMBO J. 27, 2031–2042.

28. Min, D., Kim, K., Hyeon, C., Cho, Y.H., Shin, Y.-K., and Yoon, T.-Y. (2013) Mechanical unzipping and rezipping of a single SNARE complex reveals hysteresis as a force-generating mechanism. Nature Comm. DOI: 10.1038/ncomms2692

29. Nichols, B.J., Ungermann, C., Pelham, H.R.B., Wickner, W., and Haas, A. (1997) Homotypic vacuolar fusion mediated by t- and v-SNAREs. Nature 387, 199–202.

30. Ohya, T., Miaczynska, M., Coskun, U, Lommer, B, Runge, A., Drechsel, D, Kalaidzidis, Y., and Zerial, M. (2009) Reconstitution of Rab- and SNARE-dependent membrane fusion by synthetic endosomes. Nature 459, 1091–1097.

31. Orr, A., Wickner, W., Rusin, S.F., Kettenbach, A.N., Zick, M. (2015) Yeast vacuolar HOPS, regulated by its kinase, exploits its affinities for acidic lipids and Rab:GTP for membrane binding and to catalyze tethering and fusion. Mol. Biol. Cell 26, 305–315.

32. Orr, A., Song, H., Rusin, S.F., Kettenbach, A.N., and Wickner, W. (2017) HOPS catalyzes the interdependent assembly of each vacuolar SNARE into a SNARE complex. Mol. Biol. Cell 28, 975–983.

33. Pace, N.C., and Scholtz, J.M. (1998) A helix propensity scale based on experimental studies of peptides and proteins. Biophys. J. 75, 422–427.

34. Rice, L.M., Brunger, A.T. (1999) Crystal structure of the vesicular transport protein Sec17. Mol. Cell 4, 85–95.

35. Richmond, J.E., Weimer, R.M. and Jorgensen, E.M. (2001) An open form of syntaxin bypasses the requirement for UNC-13 in vesicle priming. Nature 412, 338–341.

36. Seals, D., Eitzen, G., N., Wickner, W.T., and Price, A. (2000) A Ypt/Rab effector complex containing the Sec1 homolog Vps33p is required for homotypic vacuole fusion. Proc. Natl. Acad. Sci. USA 97, 9402–9407.

37. Schwartz, M.L. and Merz, A.J. (2009) Capture and release of partially zipped trans-SNARE complexes on intact organelles. J. Cell Biol. 185, 535–549.

38. Schwartz, M.L., Nickerson, D.P., Lobingier, B.T., Plemel, R.L., Duan, M., Angers, C.G., Zick, M., and Merz, A. (2017) Sec17 (α-SNAP) and an SM-tethering complex regulate the outcome of SNARE zippering in vitro and in vivo. eLife DOI: https://doi.org/10.7554/eLife.27396.001

39. Söllner, T., Bennett, M.K., Whiteheart, S.W., Scheller, R.H., and Rothman, J. E. (1993) A protein assembly-disassembly pathway in vitro that may correspond to sequential steps of synaptic vesicle docking, activation, and fusion. Cell 75, 409–418.

40. Song, H., Orr, A., Duan, M., Merz, A.J., and Wickner, W. (2017) Sec17/Sec18 act twice, enhancing membrane fusion and then disassembling cis-SNARE complexes. eLife DOI: 10.7554/eLife.26646.

41. Song, H. and Wickner, W. (2017) A short region upstream of the yeast vacuolar Qa-SNARE heptad-repeats promotes membrane fusion through enhanced SNARE complex assembly. Mol. Biol. Cell 28, 2282–2289.

42. Song, H. and Wickner, W. (2019) Tethering guides fusion-competent trans-SNARE assembly. Proc. Natl. Acad. Sci. USA 116, 13952–7. DOI 10.1073/pnas1907640116.

43. Song, H., Orr, A.S., Lee, M., Harner, M.E., and Wickner, W.T. (2020) HOPS recognizes each SNARE, assembling ternary *trans*-complexes for rapid fusion upon engagement with the 4th SNARE. eLife doi.org/10.7554/eLife.53559.

44. Sorensen, J.B., Wiederhold, K., Muller, E.M., Milosevic, I., Nagy, G, de Groot, B.L., Grubmuller, H., and Fasshauer, D. (2006) Sequential N-to C-terminal SNARE complex assembly drives priming and fusion of secretory vesicles. EMBO J. 25, 955–966.

45. Stepien, K.P. and Rizo, J. (2021) Synaptotagmin-1-, Munc18-1- and Munc13-1-dependent liposome fusion with a few neuronal SNAREs. Proc. Natl. Acad. Sci. USA, in press.

46. Stroupe, C., Collins, K.M., Fratti, R.A., and Wickner, W. (2006) Purification of active HOPS complex reveals its affinities for phosphoinositides and the SNARE Vam7p. EMBO J. 25, 1579–1589.

47. Stroupe, C., Hickey, C.M., Mima, J., Burfeind, A.S., and Wickner, W. (2009) Minimal membrane docking revealed by reconstitution of Rab GTPase-dependent membrane fusion from purified components. Proc. Natl. Acad. Sci. USA 106, 17626–17633.

48. Sutton, R.B., Fasshauer, D., Jahn, R., and Brunger, A.T. (1998) Crystal structure of a SNARE complex involved in synaptic exocytosis at 2.4 A resolution. Nature 395, 347–353.

49. Torng, T. and Wickner, W. (2020) Asymmetric Rab activation of vacuolar HOPS to catalyze SNARE complex assembly. Mol. Biol. Cell 31, 1060–1068.

50. Ungermann, C., Nichols, B.J., Pelham, H.R.B., and Wickner, W. (1998) A vacuolar v-t-SNARE complex, the predominant form in vivo and on isolated vacuoles, is disassembled and activated for docking and fusion. J. Cell Biol. 140, 61–69.

51. Wada, Y., Ohsumi, Y., and Anraku, Y. (1992) Genes for directing vacuole morphogenesis in Saccharomyces cerevisiae. I. Isolation and characterization of two classes of vam mutants. J. Biol. Chem. 267, 18665–18670.

52. Webb, B., and Sali, A. (2016) Comparitive protein structure modeling using MODELLER. Current Protocols in Bioinformatics. 54, 139–148.

53. Weber, T., Zemelman, B.V., McNew, J.A., Westermann, B., Gmachi, M., Parlati, F., Söllner, T.H., and Rothman, J.E. (1998) >Cell 92, 759–772.

54. White, K.I., Zhao, M., Choi, U.B., Pfuetzner, R.A., Brunger, A.T. (2018) Structural principles of SNARE complex recognition by the AAA+ protein NSF. eLife DOI: 10.7554/eLife.38888.

55. Wickner, W. (2010) Membrane fusion: Five lipids, four SNAREs, three chaperones, two nucleotides, and a Rab, all dancing in a ring on yeast vacuoles. Ann. Rev. Cell Dev. Biol. 26, 115–136.

56. Winter, U., Chen, X., and Fasshauer, D. (2009) A conserved membrane attachement site in α--SNAP facilitates N-ethylmaleimide-sensitive Factor (NSF)-driven SNARE complex disassembly. J. Biol. Chem. 284, 31817–31826.

57. Xu, H., and Wickner, W (1996) Thioredoxin is required for vacuole inheritance in *Saccharomyces cerevisiae*. J. Cell Biol. 132, 787–794.

58. Xu, H., Jun, Y, Thompson, J., Yates, J., and Wickner, W. (2010) HOPS prevents the disassembly of *trans*-SNARE complexes by Sec17p/Sec18p during membrane fusion. EMBO J. 29, 1948–1960.

59. Xu, H. and Wickner, W.T. (2012) N-terminal domain of vacuolar SNARE Vam7p promotes trans-SNARE complex assembly. Proc. Natl. Acad. Sci. USA 109, 17936–17941.

60. Zhang, Y. (2017) Energetics, kinetics, and pathway of SNARE folding and assembly revealed by optical tweezers. Protein Science 26, 1252–1265.

61. Zhao, M., Wu, S., Zhou, Q., Vivona, S., Cipriano, D.J., Cheng, Y., and Brunger, A.T. (2015) Mechanistic insights into the recycling machine of the SNARE complex. Nature 518, 61–67. DOI 10.1038/nature14148.

62. Zick, M. and Wickner, W. (2013) The tethering complex HOPS catalyzes assembly of the soluble SNARE Vam7 into fusogenic *trans*-SNARE complexes. Mol. Biol. Cell 24, 3746–3753.

63. Zick, M., and Wickner, W. (2014) A distinct tethering step is vital for vacuole membrane fusion. eLife DOI:10.7554/eLife.03251.

64. Zick, M. and Wickner, W. (2016) Improved reconstitution of yeast vacuole fusion with physiological SNARE concentrations reveals an asymmetric Rab(GTP) requirement. Mol. Biol. Cell 27, 2590–2597.

65. Zick, M., Orr, A., Schwartz, M.L., Merz, A.J., and Wickner, W.T. (2015) Sec17 can trigger fusion of trans-SNARE paired membranes without Sec18. Proc. Natl. Acad. Sci. USA, 112, E2290–E2297.

